# High-Resolution Laminar Identification in Macaque Primary Visual Cortex Using Neuropixels Probes

**DOI:** 10.1101/2024.01.23.576944

**Authors:** Li A. Zhang, Peichao Li, Edward M. Callaway

## Abstract

Laminar electrode arrays allow simultaneous recording of activity of many cortical neurons and assignment to layers using current source density (CSD) analyses. Electrode arrays with 100-micron contact spacing have been used to estimate borders between layer 4 versus superficial or deep layers, but in macaque primary visual cortex (V1) there are far more layers, such as 4A which is only 50-100 microns thick. Neuropixels electrode arrays have 20-micron spacing, and thus could potentially discern thinner layers and more precisely identify laminar borders. Here we show that laminar distributions of CSDs lack consistency and the spatial resolution required for thin layers and accurate layer boundaries. To take full advantage of high density Neuropixels arrays, we have developed approaches based on higher resolution electrical signals and analyses, including spike waveforms and spatial spread, unit density, high-frequency action potential (AP) power spectrum, temporal power change, and coherence spectrum, that afford far higher resolution of laminar distinctions, including the ability to precisely detect the borders of even the thinnest layers of V1.

## Main Text

The laminar organization of the neocortex reflects the distributions of cells with different soma shapes and sizes, packing densities, and patterns of synaptic inputs and axonal outputs (*1, 2*) (Figure 1A). To link functional properties to cell types and circuits it is useful to identify the laminar locations of recorded cells. Historically this was first done by using a single electrode and marking recording locations with periodic lesions that are typically rather large and asymmetric, leaving considerable uncertainty. In the context of visual cortical circuits and function, the visual receptive fields of neurons have often been characterized with respect to laminar depth (*3–5*).

**Figure 1.**
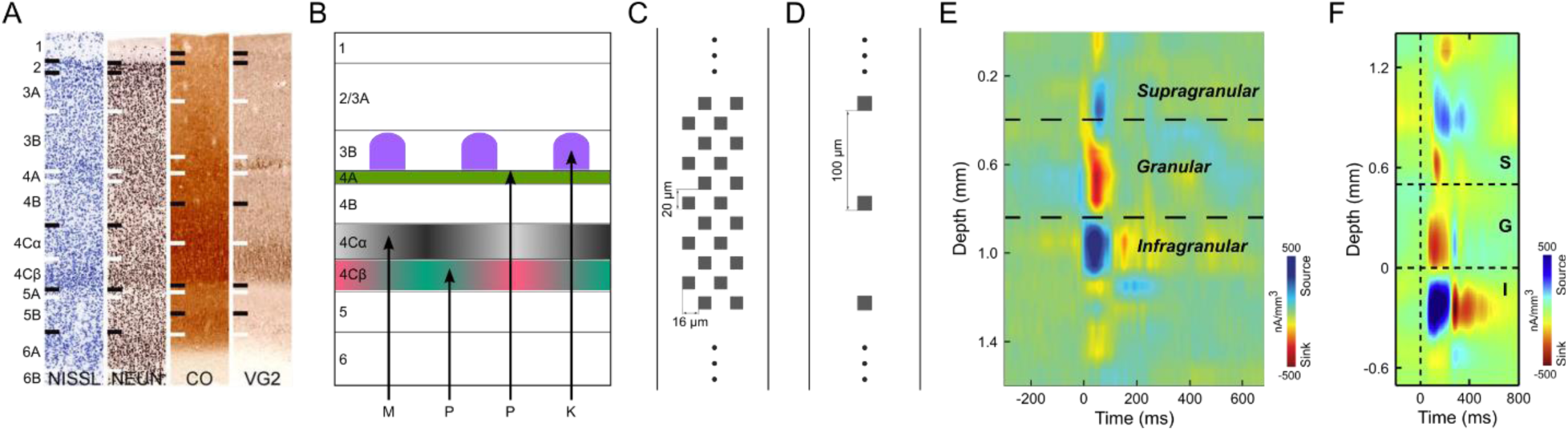
Cortical layers of V1. (**A**) Brodmann’s laminar divisions of macaque monkey V1 revealed with different histological staining methods (Nissl for cell density, Anti-NeuN for neuronal density, CO for metabolic activity and VG2 for glutamatergic axon terminals from thalamus). Adapted from (*1*). (**B**) Schematic diagram of V1 layers that receive different functional/anatomical types of LGN afferent input (*13*). LGN magnocellular (*M; L+M achromatic, grey in 4Cα*), parvocellular (*P*: *L/M cone-opponent, red-green in layer 4Cβ; S-OFF, lime in layer 4A*) and koniocellular (*K: S-ON, purple in layer 3B CO blobs*). (**C**) Electrode spacing and layout of Neuropixels 1.0 probe. (**D**) Layout of probes that were used in E and F. (**E**) An example of V1 CSD profile in awake recordings using a 16-channel linear probe (*8*). (**F**) Another example of V1 CSD profile in anesthetized recordings using a 24-channel linear probe (*9*). Supragranular (S), granular (G) and infragranular (I) layers were delineated in E and F mainly based on transitions between the short-latency granular layer sink and adjacent sources.

The development of laminar electrode arrays has allowed simultaneous recordings from many locations across cortex enabling current source density (CSD) analysis (*6, 7*) based on Local Field Potentials (LFP) in response to full-field flashed visual stimuli. The estimated laminar boundaries from CSD profiles allow direct assignment of units recorded on particular electrode contacts to their cortical layers (*8–10*). Typically, a fast current sink in the CSD profile following a full-field dark to light visual stimulus transition is attributed to the lateral geniculate nucleus (LGN) thalamic-recipient granular layer (L4) and the top and bottom of this sink are used to specify supragranular and infragranular divisions. The recent advent of long and very high-density Neuropixels arrays (*11*) has the theoretical potential to allow the identification of more layers across greater cortical depths and higher resolution identification of laminar boundaries. This potential could be particularly impactful for laminar identification in the primary visual cortex (V1) of monkeys where V1 is composed of multiple thin and functionally disparate sublayers, including at least 5 distinct layers receiving input from functionally different LGN afferent types (*12, 13*). Neurons in L4Cβ of V1 have small cell bodies and receive input from LGN parvocellular L/M cone-opponent afferents that terminate in 4Cβ, whereas neurons in 4Cα have slightly larger cell bodies and receive LGN magnocellular achromatic inputs that terminate only in 4Cα. The thin layer 4A (L4A) is packed with small granule cells that receive parvocellular S-cone OFF (S-OFF) inputs, and layer 3B (L3B) cells in high Cytochrome Oxidase (CO) intensity blobs receive LGN koniocellular S-cone ON (S-ON) projections (Figure 1B). Layer 6 (L6) also receives direct LGN input and can be divided into upper (L6A) and lower (L6B) halves based on the higher density of cell bodies in 6A (*12, 14*).

Previous use of laminar electrode arrays and CSD analyses for laminar assignment in monkey V1 (*8–10*) is limited (Figure 1E, F), because the vertical spacing of electrodes is too large (e.g. 100μm) to precisely assign laminar boundaries or to detect thin layers such as L4A, which is less than 100 µm thick. Moreover, with sparse electrode contacts, only a few single units are isolated from layers such as 4Cβ of V1, which are composed of very small neurons (∼5-8µm in diameter). In this study, Neuropixels probes (*11*), with 384 simultaneously active recording channels and 20µm vertical spacing spanning 3.84mm were used to record representative samples of neurons across the full depth of V1 (Figure 1C). A prior study used CSD analysis with Neuropixels electrodes for laminar identification in V1 of macaque monkeys (*15*), but did not attempt to take advantage of the potential for high-resolution laminar assignment; instead, recordings were averaged across four contacts, smoothed (σ=120μm), and laminar boundaries approximated based on expected depth from a single CSD transition attributed to the granular/infragranular border. It might be expected that high-resolution CSD profiles recorded using Neuropixels with 20µm vertical electrode spacing can provide finer details to identify more V1 layers with higher accuracy. In the present study, however, we show that CSD profiles are low resolution and inconsistent across both penetrations and visual stimulus conditions, making it impossible to summarize a stereotypical CSD profile that can be used to reliably and accurately relate electrode contacts to distinct layers and sublayers in V1.

In physiological situations, cortical LFPs arise mainly from synchronized synaptic activities (*16*). Because synapses are widely distributed on dendrites, the current sinks and current sources of synaptic events revealed by CSD can be far from the cell body (*17*). By convention, anatomical properties, such as neuron density (*18*) and soma size/shape are the main metrics used for anatomical laminar structure identification. Therefore, the boundaries between sinks and sources of CSD profiles do not necessarily correspond closely to laminar borders using anatomical features. Here we describe the use of alternatives to CSD profiles for linking electrical activity to layers, with the key point being the use of signals with higher-spatial resolution than CSDs and potentially having a better match to layers identified with cell body sizes, and cell densities in histological staining. High frequency (300-3000Hz) Action Potential (AP) band signals are related to the combined voltage fluctuations resulting from neuronal action potential firing, with the largest amplitude signals originating at cell bodies. These high-frequency signals have much faster attenuation across space compared to LFPs (*17*), potentially allowing for higher spatial resolution. Thus, we formulated several metrics based on AP band signals and found that they show various degrees of similarity to the laminar distributions of neuron density and soma size/shape across cortical layers. Since some layers share similar anatomical properties and are hard to differentiate even in histological staining, we also deployed colored visual stimuli that preferentially activate LGN afferents terminating in specific layers (Figure 1B). By combining selected identification metrics under different stimulus conditions, we show how high-resolution laminar structures of V1, including major layers and sublayers, can be consistently and accurately identified *in vivo* without the assistance of postmortem histological staining.

## Results

35 vertical electrode penetrations (at least 1mm apart, typically far more, see Figure S11) were made using Neuropixels 1.0 probes in primary visual cortex (V1) of two anesthetized/paralyzed rhesus macaque monkeys (17 penetrations in monkey A and 18 in monkey B), of which 31 are included in our analyses (See Methods for criteria used to exclude penetrations A7, A8, A17 and B18). Of these 31 penetrations, 20 were centered on Cone Opponent Functional Domains (COFD) and 9 on achromatic ON/OFF domains revealed by intrinsic signal imaging (ISI) (*19*). The remaining 2 penetrations were targeted to regions outside these domains. ISI was also used to reveal ocular dominance columns in V1 and the border between primary and secondary (V2) visual cortex. We will first present results using CSD analysis demonstrating its limitations in spatial resolution and consistency, and then introduce new metrics based on AP band signals to effectively distinguish between even thin V1 layers and to precisely define laminar boundaries. We then describe a set of steps for applying these metrics to guide identification of all laminar boundaries in V1. Note that the labeled layer boundaries in all figures, including those illustrating CSD data, were determined using the full set of metrics; therefore these plots illustrate borders that might not be apparent solely from visualization of the analyses being illustrated in a particular plot. Discrepancies between labeled layer boundaries and borders revealed in plots based on particular metrics highlight the limitations of those metrics, as is most clear from the CSD analyses described below.

### CSD profiles are low-resolution and inconsistent for the same stimulus

The shortest latency current sink in cortical area V1 CSD profiles evoked from flashed, full-field visual stimuli is thought to correspond roughly to L4C, which receives the most dense LGN inputs; and the transition from this sink to a deeper current source has been used to identify the border between granular L4 and infragranular L5 (*8–10, 15*). CSD profiles (Figure 1E, F) in many previous studies were obtained using 16, 24, or 32-channel linear probes with 100µm vertical spacing (Figure 1D). The signal detection gap in these electrodes precludes identification of thin layers or sub-layers such as 4A, 4Cα and 4Cβ. Interpolations were usually applied to these CSD profiles so that layer borders could be drawn at sink-source boundaries at a resolution higher than 100µm, however, the location of a sink-source boundary might deviate substantially from the same boundary in CSD profiles produced by Neuropixels, which have a 20µm vertical spacing. Figure 2A shows an example 20µm spacing CSD profile obtained using Neuropixels. At this resolution the CSD signals are highly fragmented due to more or less locally balanced high resolution sink-source patterns along the probe. While there appear to be alternating contacts with a strong short latency sink within L4C, these only span a depth of about 200µm, far narrower than L4C, leaving considerable uncertainty about where boundaries should be drawn. To simulate results with lower resolution arrays and to potentially facilitate layer identification by eliminating local current spread, we applied gradually increasing width (σ=20, 30, 40µm) gaussian filters to derive sink-source patterns that more closely matched the scale of most cortical layers. In figures 2B-D, it can be seen that, overall, CSD profiles become more stable with increasing degrees of smoothing while missing small details. This gaussian filtering approach yielded nearly equivalent results to decreasing spatial resolution by calculating CSDs using down-sampling to skip 1, 2, 3, or 4 contacts, which mimicked the CSDs that would have been generated by linear probes with 40, 60, 80, or 100µm spacing (Figure S1). To better match the scale of cortical layers while keeping as much detail as possible, all subsequent CSD analysis results illustrated were gaussian filtered with σ=30µm, which was almost identical to CSD results from analyses with down-sampling at ∼60µm contact spacing.

**Figure 2.**
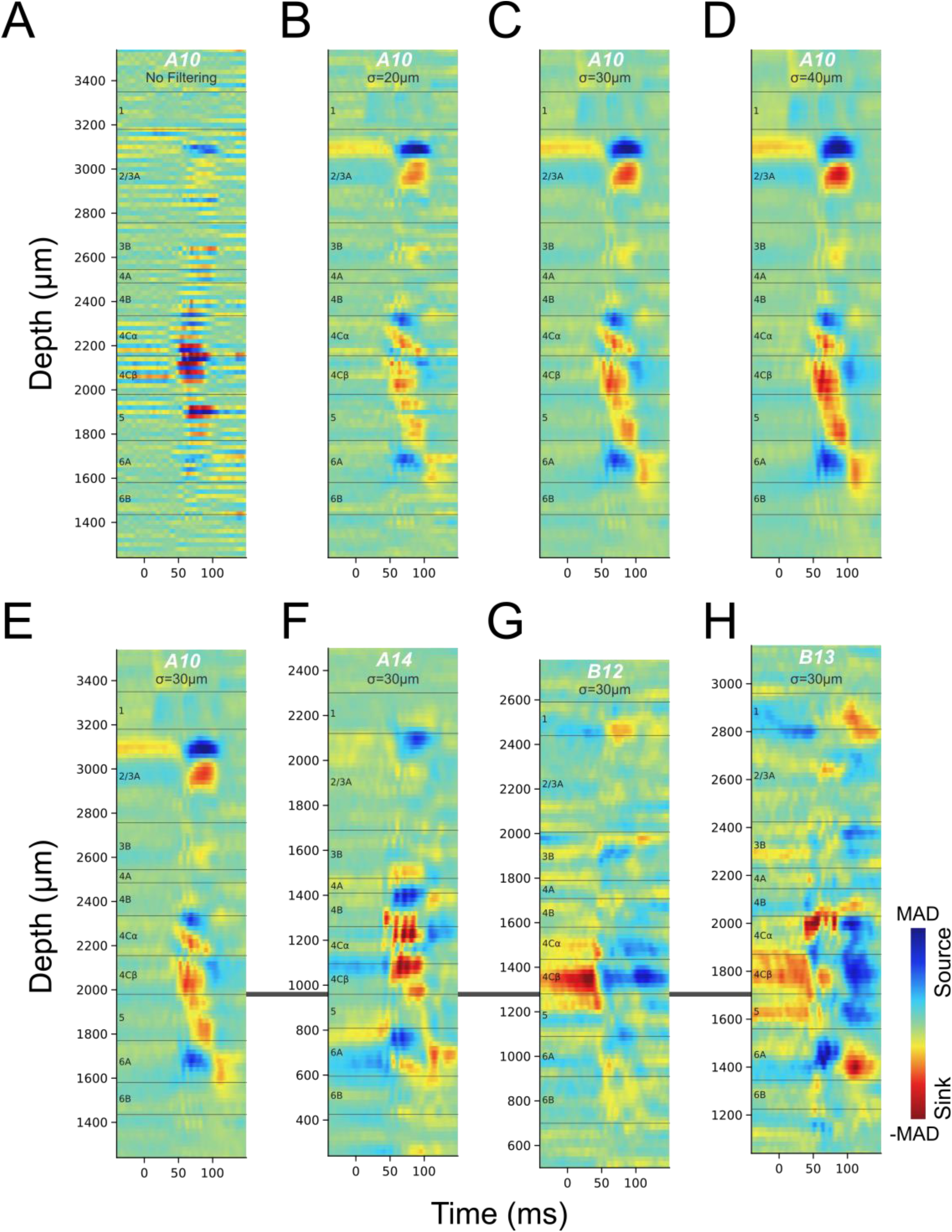
CSDs evoked by a black to white visual stimulus transition through the dominant eye, all targeting achromatic ON domains. The layer boundaries drawn here were determined using the methods and data described in the subsequent sections. (**A**) CSD profile from the LFP traces of all contacts with 20µm depth spacing (Penetration A10). (**B-D**) Gaussian filtering of the CSD profile in A along the depth of probe with σ equals 20µm (B), 30µm (C) or 40µm (D). (**E-H**) CSD profiles from four penetrations (E: A10, F: A14, G: B12, H: B13) generated using 30µm gaussian smoothing. The CSD profiles were aligned relative to the border between L4Cβ and L5, and CSD scales were each independently color-mapped relative to the maximum absolute deviation (MAD) in each profile.

The most used visual stimulus in past studies is a full screen switching from black to white. We therefore first compare our spatially smoothed CSD profiles using black to white transitions with the low spatial resolution CSD profiles (Figure 1E, F) reported in previous studies (*8, 9*) in which a stimulus presentation screen was also flipped from black to white for ∼100ms. Figures 2E-H show CSD profiles from four penetrations generated by flashing from a black to a white screen through the dominant eye (black to white every 3 seconds); these CSDs clearly lack consistency across penetrations. The criteria from previous studies using the best boundary between an initial sink and a deeper source to mark the border between L4C and L5 is not tenable because sometimes no such transition is apparent and there is never a clearly identifiable transition. More importantly, the observed transitions do not consistently match to layer identification using the methods that we establish and validate below. For example, in Figure 2E, criteria applied in previous studies would mark the bottom of L4C at 1800µm, where the initial sink transitions to a source (compare Figure 1E to 2E). But this border location is highly unlikely because there would be insufficient space to accommodate the known thickness of L5 and L6 before our unambiguous identification of the L6/white matter border (see below). Furthermore, in some cases the CSD profile in response to a black to white screen has a zero latency sink in L4C that transitions to a current source (Figures 2G, H). A prior study has shown that the zero latency sink corresponds to residual signal from the prior white to black transition (*20*) which occurs 1.5 seconds earlier in our experiments. We note that this signal is much stronger in layer 4Cβ than 4Cα, consistent with more sustained activity in parvocellular LGN neurons and an asymmetry in which there is more sustained activity in the dark (black stimulus) condition.

### Inconsistency and stimulus dependency of CSD profiles

Here we explore the consistency of CSD profiles both across individual penetrations and across visual stimulus conditions. In view of the ambiguity in CSD profiles that might persist beyond the time before a new stimulus transition (Figure 2G, H), we use ΔCSD (See Methods) in subsequent CSD analyses to allow comparisons of CSDs under additional stimulus conditions and to further assess variability in CSD responses. In addition to conventional black and white stimuli, we constructed L/M cone-opponent colors and S/(L+M) cone-opponent colors, to preferentially activate cone-opponent parvocellular LGN inputs to L4Cβ (L/M opponent) or L4A (S-OFF), or koniocellular LGN inputs (S-ON) to L3B blobs (Figure 1B). Figure 3 shows ΔCSD profiles generated in response to various stimulus conditions for three penetrations. Within each penetration, different stimulus conditions usually evoked different ΔCSD profiles, some of which could differ drastically, e.g. black vs. white (Figure 3C), or dominant eye vs. non-dominant eye (Figure 3B). Meanwhile, ΔCSD profiles generated using the same color or ocular dominance also show various degrees of inconsistency across penetrations. The most consistent and clear generation of a fast sink corresponding to L4C is observed with a black stimulus (white to black) presented to the dominant eye, whereas the response to a white stimulus is more variable and can evoke an early L4C source in some penetrations (Figure S2, S3). Note that even in these best-case conditions, the L4C sinks leave considerable uncertainty about where laminar boundaries should be drawn.

**Figure 3.**
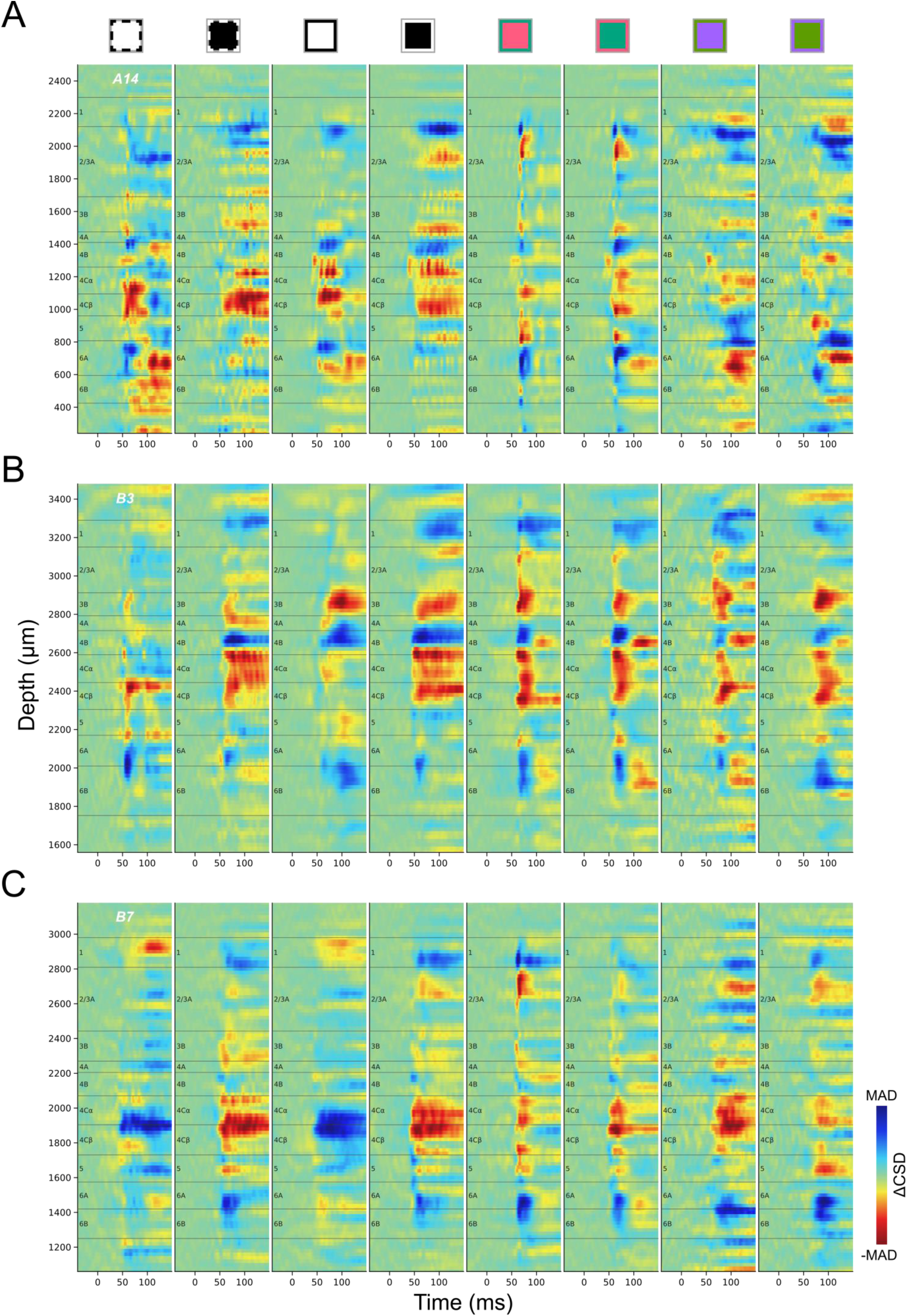
ΔCSD profiles of three penetrations (A: A14, B: B3, C: B7) under different stimulus presentations. Square icons above CSD profiles indicate the different color transitions used to generate the CSDs, as well as which eye was stimulated. The outline of each square indicates the color of the screen before the stimulus transition at time zero, while the filled color indicates the flashed color (Achromatic, Black and White; or color pairs in DKL color space; See Method). Black and White correspond to achromatic stimuli; Red to L-ON, M-OFF; Green to M-ON, L-OFF; Purple to S-ON, (L+M)-OFF; and Lime to S-OFF, (L+M)-ON. Dashed contours (two leftmost columns) indicate visual presentation to the non-dominant eye and solid contours indicate stimuli presented to the dominant eye. Color maps for ΔCSD profiles were independently scaled according to the maximum absolute deviation (MAD) in each profile.

Responses to L/M cone-opponent colors also differ from black/white responses in that they often lack a current source below L4C that could help to identify the L4C/L5 border (Figure 3). The variability of ΔCSD profiles across penetrations makes it difficult to summarize a single stereotypical profile that can be used as a template to delineate layer boundaries for each penetration. Even using the most stable stimulus, black screen to the dominant eye, landmarks are often missing that would be needed to reliably identify major layers and sublayers. These results suggest that new methods are required to accurately and consistently identify most of the known layers and sublayers.

#### Action potential-based metrics for laminar identification

In the following sections we describe several metrics extracted from high frequency action potential (AP) band signals that can be informative for more definitively identifying the precise location of laminar borders and for identifying borders that are not typically seen with conventional CSD analyses. These metrics can be categorized into two groups. Metrics in the first group were features derived from the density and spike shapes (“waveforms”) of units identified through spike sorting. These were independent of the visual stimuli used and reflected the inherent properties of spikes and the spatial relationships between locations of spike initiation/propagation and electrode contacts. Metrics in the second group were derived directly from AP signal responses to full-field flashed visual stimuli. We also compared these measures for responses to different visual stimulus conditions designed to potentially reveal locations receiving input from different types of LGN afferents.

### Unit Density

The neuron packing density is a clear anatomical property that is useful for differentiating cortical layers with histological staining. Neuron density depth distributions derived by counting immunolabeled neurons in confocal microscopy (*18*) clearly show packing density patterns that corresponded to each layer. In macaque monkey V1, L4Cβ has the highest packing density. Moving downward, there is a density trough in L5 and then an increase again in L6. Moving upward from L4Cβ, density decreases across L4Cα and L4B, until it rises to a small peak that corresponds to the thin L4A. To estimate neuron density from our electrical recordings we can plot the density of units resolved during spike sorting. The accuracy of this estimation depends, of course, on the ability to consistently resolve large numbers of units across the depth of cortex and is also subject to potential sampling biases that might make some spike waveforms easier to isolate than others. Figure 4A plots unit density (See Methods) for an example penetration across V1 and into the white matter (WM), and shows clear changes with depth that partially capture the shape of the expected neuron density distribution and provide landmarks helpful for inferring the likely identities of some layers. The obvious peak around the electrode depth at 2400µm (distance relative to deepest active electrode contact) in Figure 4A corresponds to L4C and the trough below that peak is expected from the low neuron density in L5. Below L5, the expected transition to higher density in L6A is observed, as well as decreased density in L6B. These same transitions can be observed in the histological profiles of neuron density described in figure 6 of the study from Kelly and Hawken (*18*); note however that the electrical signals contrast strongly with the histological density at the L6/WM border because high densities of units isolated from axons can be detected in the white matter. As shown below, these can be distinguished by their unique spike waveforms.

**Figure 4.**
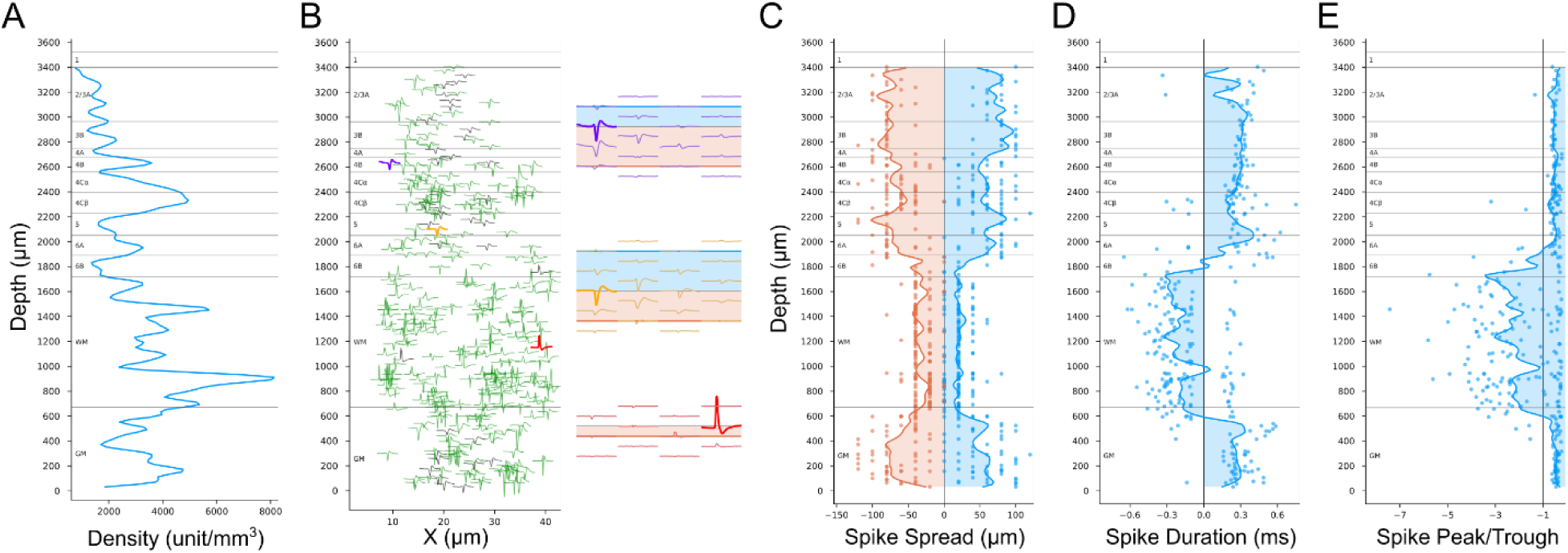
Unity density and spike waveforms features across cortical depth. (**A**) Unit density distribution along the probe for one penetration (A6). (**B**) Average spike waveform of maximum height was shown for single unit (green) or multi-unit (gray). The insets show the spike spatial spread on the Neuropixels probe for the units with corresponding color, and the filled bands indicate the upward spread (blue) and downward spread (orange). (**C**) Upward (blue) and downward (orange) spike spread. (**D**) Spike duration. (**E**) Spike peak-trough amplitude ratios. A-E are from the same penetration (A6). In C-E, each dot represents a unit, and lines are smoothed distributions along the probe (See Methods). The transitions to small spike spread in C, sign reversals of spike duration in D and crosses at −1 of peak-trough ratio in E, are strong indicators for the locations of the border between white matter (WM) and gray matter (GM).

### Metrics from Spike Waveforms

Cell body size and shape are additional anatomical features related to laminar changes in histological staining. Densely packed neurons usually have small cell bodies, e.g., the majority of neurons in L4Cβ and L4A are small granule neurons. Conversely, sparsely packed neurons in L4B and L5 are mostly large pyramidal or spiny stellate neurons. The spiking signal of a neuron propagates through extracellular space with decaying amplitude. The travel distance before it is attenuated to the noise level can be measured in a group of electrodes of which the spacing is less than the decay distance (Figure 4B insets). The spike spatial spread measured on electrodes may be related to soma size/shape. Assuming two neurons are located at the same horizontal distance from the probe, then the one with the larger soma, or a more elongated shape along the axis of the probe should be detected on more electrode contacts, resulting in larger spike spread.

Figure 4C plots the distributions of spike spread distances measured for units isolated on contacts at different laminar depths. Although the distributions are somewhat granular owing to the fixed 20μm spacing between electrode contacts, the smoothed mean distribution is clearly at its lowest in the white matter, and within the cortex the lowest mean values are observed across L4C. There is also an increase in spread from L4C to L5 as expected from the spikes from larger L5 cells. While other transitions from this measure do not always have a close match to the boundaries identified with other measures, spike spread does prove to be particularly useful to differentiate white matter from gray matter because the spread of axonal spikes or noise detected in white matter is much smaller than that of spikes from cell bodies. The location of the L6/WM border is further corroborated by the expected transition to inverted spikes that are measured from axons in the white matter (Figure 4B, compare red inverted spikes to conventional purple and orange spikes). These inverted spikes have a negative spike duration (difference in time from peak to trough) and a large, negative peak-trough ratio (< −1). Depth plots of spike duration (Figure 4D) and peak-trough ratio (Figure 4E) show clear and sharp transitions at the L6/WM border. In these examples, there is an additional transition as the electrode penetrates across the white matter and back into L6 of the V2 cortex that is folded under the opercular cortical surface. Note that a few inverted spikes are also detected in a L6B transition zone where cell bodies are intermingled with a larger density of axons than in more superficial layers, as well as in L4Cβ where parvocellular LGN afferents are very dense and their spikes can be isolated relative to the surrounding small spikes from 4Cβ spiny stellate neurons.

Unit density, spike spread, spike duration and spike peak-trough ratio are all stimulus-independent spike features, and together can be used to identify the approximate locations of L4C and L5, as well as the L6/WM border. These are crucial landmarks that can serve as anchors for the validation and assignment of the identities of additional borders and layers derived from other features. Use of this group of metrics for laminar assignment depends on numbers of detected and resolved units; thus, probes with high contact numbers and densities, such as Neuropixels, are essential. In practice, cortical health is also important as damage or conditions reducing the number of active and resolvable spikes will drastically reduce the number of detectable units, and the spike sorting process may exclude small amplitude spikes or unresolvable spikes further reducing the yield, in turn impairing the accuracy of estimating neuron density and soma size/shape through these metrics. We next explore a second group of complementary metrics derived directly from stimulus-evoked AP band signals that may resolve additional laminar boundaries and/or display sharper transitions at laminar borders. These measures also circumvent limitations imposed by numbers of units detected and resolved and might prove useful with lower density electrode arrays.

### Power Spectrum of AP Signals

Below we introduce three additional AP-band metrics that are all derived from the high-frequency voltage fluctuations measured in the AP band (300-3000Hz). Two of these metrics, “AP Power” and “Coherence”, each generate spectral plots based on voltages measured across cortical depth, plotted against the frequencies of the voltage fluctuations. Because these plots show the average values during a response window (10-150ms after the visual stimulus transition), they are insensitive to temporal differences in responses that might be expected due to different response latencies at different depths, such as faster transitions expected at LGN recipient layers or layers receiving shorter latency LGN inputs from the faster propagating magnocellular afferents that terminate in L4Cα.

In cortical tissue, AP band voltage fluctuations result mainly from the superposition of neuronal spikes, which typically feature a trough-peak deflection of extracellular potentials (Figure 4B purple and orange). There are several properties of spiking activity evoked by visual stimuli that are expected to influence laminar power spectrum profiles. First, there is expected to be an overall influence of neuron density. For example, if every spike were to have the same shape and amplitude and there were no waveform cancelling due to temporal overlap, then the Root Mean Square (RMS) amplitude of the AP signal on a recording channel would be directly proportional to number of spikes. This factor might be expected to result in some similarity between laminar power profile and that of neuronal density, given the firing rates of neurons across layers are also similar. Second, the number of spikes detected is also expected to reflect the degree to which neurons close to each contact are activated by the particular visual stimulus that preceded the response window. Since all visual stimuli used were full-field and not spatially structured, this might be expected to result in a strong relationship to layers that respond to full-field stimuli, as well as those receiving direct input from LGN axons responding to these stimuli and whose APs also contribute to measured voltage changes. Taken together, it is possible that the firing rate profile across depth could counteract the depth profile of neuron density, and the resulting depth profile of power could deviate far from the depth profile of neuron density. Nevertheless, we found that depth profile of firing rate to these full-field stimuli roughly matches the depth profile of neuron density (Figure S4B, C), making power spectrum profile a useful estimator of neuron density.

Figure 5 plots RMS^2^ Power Spectra (*P*_*f*_, See Methods) during a response window from 10-150ms after various stimuli, averaged across trials. In the majority of penetrations, laminar power spectra plots have at least two clear laminar bands corresponding to L4C and L6A. Under some stimulus conditions an additional thin band is seen in L4A. These signatures almost always precisely align with sharp transitions at the top and bottom of L5, marking the L4Cβ/L5 border and the L5/L6 border. In most cases the top of the L4C band corresponds to the L4B/L4C border. But in some cases (e.g. Figure 5B) the transition to lower power occurs below the expected L4B/L4C border as identified from other measures from the same penetration, such as response latencies and the known thickness of L4Cα relative to L4Cβ (see below). In cases where a thin L4A band is visible, the borders can precisely identify the top and bottom of that layer. These patterns match with the expectation of lower AP power in layers with low neuronal density that do not receive direct LGN input (e.g. L4B and L5). Insofar as both L6A and L6B receive direct LGN inputs (*14*), the greater AP power in 6A is likely due to higher neuron density. The stronger signal that is sometimes observed in L4Cβ relative to L4Cα might also be expected from the higher neuronal density in L4Cβ, but this is not a consistent feature of laminar power spectra. It is also possible that differential patterns under various stimulus conditions might relate to the columnar organization or laminar organization of cone-opponent mechanisms in V1. For example, S-OFF LGN inputs to L4A might be expected to result in consistent L4A activation to the S-OFF stimulus, but we do not find highly predictable relationships between the particular-colored stimuli that differentially activate layers to the ISI-identified cone-opponent columns targeted by the electrode penetrations. Nevertheless, the use of multiple stimuli appears to increase the chances of detecting these unpredictable activation patterns.

**Figure 5.**
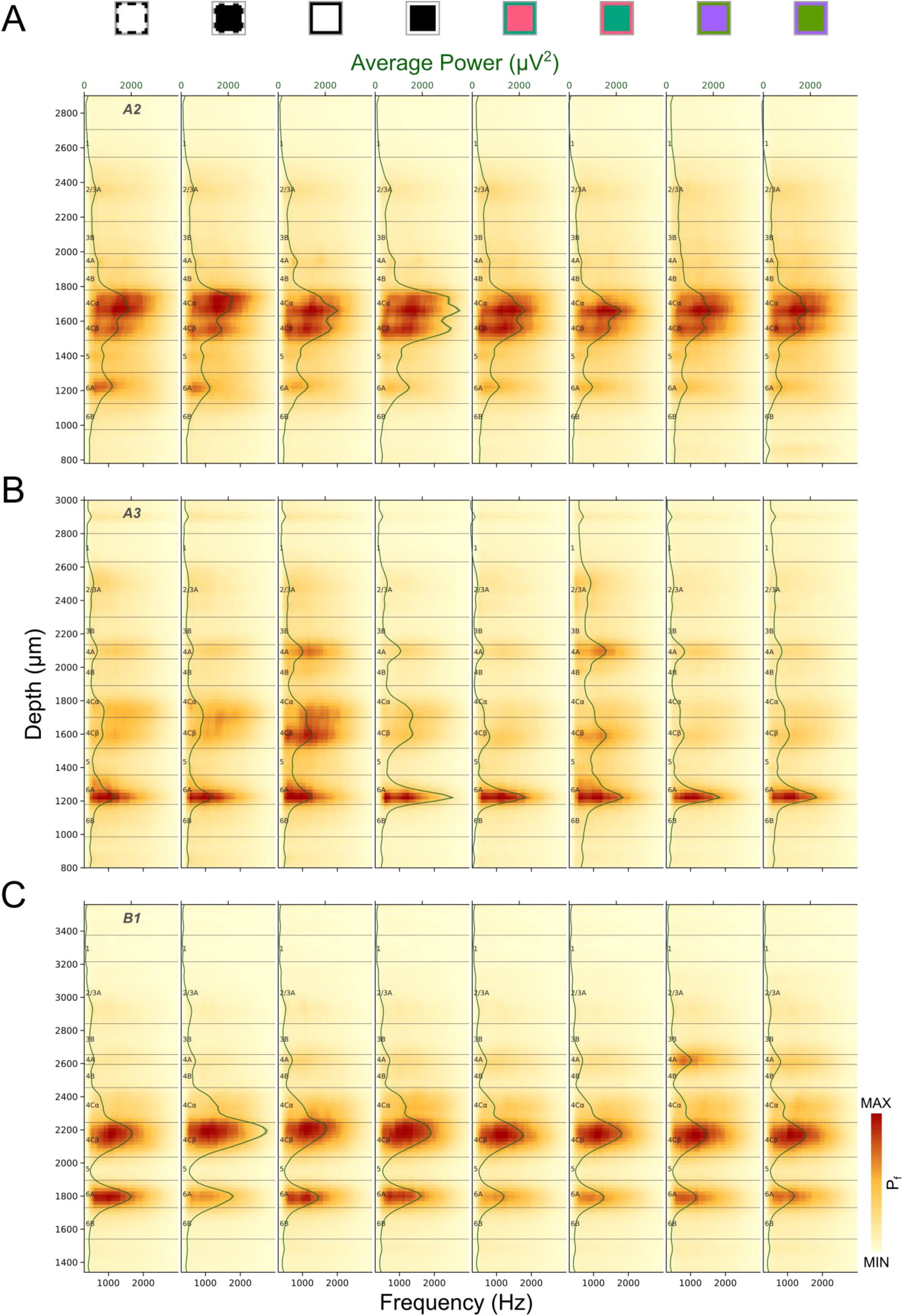
Laminar power spectrum (P_f_) profiles for three penetrations (A: A2, B: A3, C: B1) generated in response to different stimulus conditions. Power spectrum profiles were independently color-mapped according to the minimum and maximum power in each profile (See Figure 3 legend for stimulus conditions corresponding to icons above plots). The green lines and corresponding shaded ribbons show the Mean±SEM power across frequency (300-3000Hz). Note that SEM is roughly the same as the thickness of the line.

### Local Coherence spectrum of AP Signals

While overall AP power, as described above, clearly generates signals that are reliable and precisely related to cortical layers, differential AP power on different electrode contacts might sometimes result from stochastic variability in the precise locations of electrode contacts in relation to neuronal cell bodies. For example, the AP amplitude measured from a particular neuron is typically much larger on one contact than other nearby contacts (Figure 4B insets). These differences might contribute significantly to differential power signals, suggesting that it might be useful to compare power to an alternative measure that is independent from spike amplitudes. We calculated coherence spectra (*C*_*f*_, See Methods) between recording channels to generate a measure that is independent from variation due to different spike amplitudes (Figure S4A). Coherence represents normalized phase relationships between two signals, where zero coherence indicates random phase relationships and one indicates deterministic phase relationships. As seen in Figure 4B, spikes from one neuron usually spread to a group of nearby recording contacts with very small time delays and much attenuated amplitudes. These relationships are expected to result in significant local coherence between neighboring channels. Assuming spikes of a neuron spread to at least two adjacent recording channels, then the coherence between the two channels would be directly proportional to number of spikes, independent of spike amplitude. Thus, coherence is expected to be related to power in that both are related to the number of action potentials that are evoked around an electrode contact, but they differ in that coherence is insensitive to spike amplitudes.

Figure 6 illustrates laminar patterns of coherence spectra during responses to various visual stimuli for 3 electrode penetrations. Like laminar AP power spectra, coherence is generally greatest in L4C and L6A. While there are generally fewer and less distinct laminar transitions for coherence spectra than for power spectra, there are nevertheless sometimes cases where they can resolve uncertainties. In many penetrations coherence is also a useful and clear indicator of the white matter location due to the transition to a minimal value at the L6/WM border.

**Figure 6.**
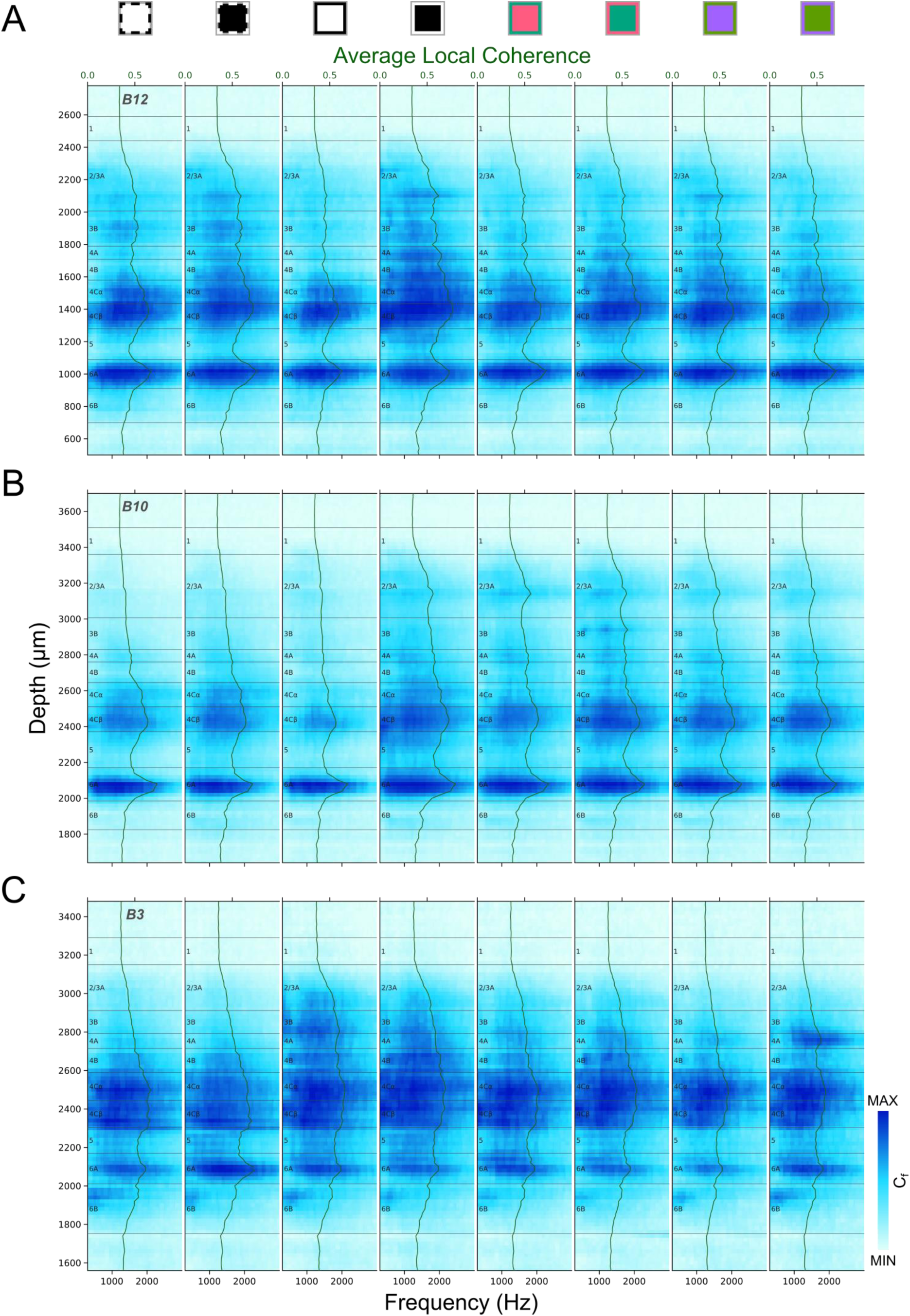
Local coherence spectrum (C_f_) profiles for three penetrations (A: B12, B: B10, C: B3) generated in response to different stimulus conditions. Local coherence profiles were independently color-mapped according to the minimum and maximum local coherence in each profile (See Figure 3 legend for stimulus conditions corresponding to icons above plots). The green lines and corresponding shaded ribbons show the Mean±SEM local coherence across AP frequency (300-3000Hz). Note that SEM is roughly the same as the thickness of the line.

### AP Power temporal change profile

Another measure that is expected to be closely related to both neuronal density and responsiveness is the AP power change relative to average power in the baseline time window (ΔP/P, See Methods). Because this measure subtracts baseline power it is expected to be less sensitive than AP power spectra to variations in spike amplitude. This measure also has the advantage of revealing timing between layers that might result from different response latencies. It can be seen in Figure 7 that responses in L4Cα are typically faster than L4Cβ, consistent with previous findings that the magnocellular LGN afferents and their recipient neurons in L4Cα have shorter response delays than for the parvocellular recipient neurons in L4Cβ (*13, 21*). Different colored visual stimuli also often generate distinctly different laminar and temporal patterns corresponding to L4A, L4Cα, or L4Cβ, that also vary between penetrations in their latency and response strength (Figure S2, S3). For example, for each of the three penetrations shown in Figure 7, the S-OFF parvocellular afferent-recipient L4A is clearly defined by high ΔP/P with longer latency than for L4Cα, in response to at least one of the colored visual stimuli. But for one penetration (Figure 7A) the L4A band is clearest in response to the S-ON stimulus, while for another (Figure 7C) it is only observed in response to the S-OFF stimulus. There is also variability between penetrations and stimuli in the temporal sequence observed for activation of the LGN-recipient layers L4A, L4Cα, L4Cβ and L6A. This is particularly striking for the responses to the S-ON stimulus for the penetration shown in Figure 7B, where there are distinct transient increases that clearly and sharply define the borders of each of these layers and which occur first in L4Cα, then L6A, followed by L4A, and finally L4Cβ. More typically, response latencies are similar in L4A and L4Cβ. In all penetrations the responses in L4Cα have the shortest latency. Overall, ΔP/P profiles often provide laminar information that is more difficult to deduce from other measures, particularly for L4Cα, L4Cβ, or L4A.

**Figure 7.**
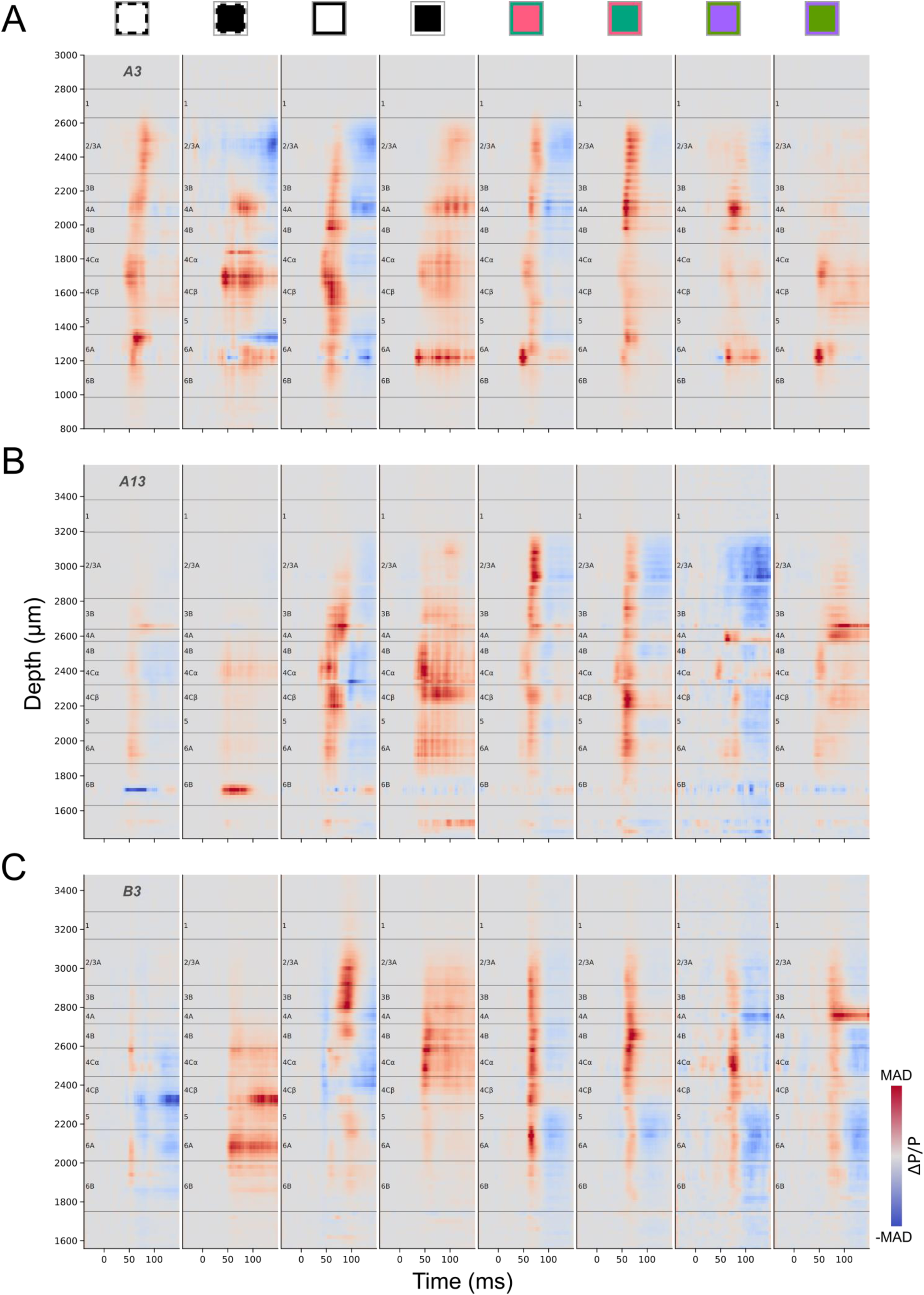
ΔP/P profiles for three penetrations (A: A3, B: A13, C: B3) generated in response to different stimulus conditions. ΔP/P (See Methods) profiles were independently color-mapped according to the maximum positive (red) and negative (blue) absolute deviation (MAD) for each profile (See Figure 3 legend for stimulus conditions corresponding to icons above plots).

#### Combining Metrics for Accurate, High Resolution Laminar Delineation

While many of the metrics described above consistently and precisely identify some laminar boundaries, there is not a single metric that can identify all layers. For example, some metrics are most closely related to anatomical properties such as neuron density or soma size, while others reveal differences related to functional properties, such as visual stimulus preference or response latency. In addition, some metrics are very good for providing anchor points, such as the general location of a layer, but are not as good as others for defining precise borders. Finally, the data also demonstrate that there is considerable variability between penetrations, such that the metrics and visual stimuli that best reveal laminar boundaries differ between penetrations. Below, we therefore outline a general order in which various criteria might be ideally applied to accurately identify as many laminar boundaries as possible for a given penetration, but common sense should be used to estimate confidence levels for any particular set of data based on available data quality and consistency.

Below and in Figure 8 we illustrate a general workflow for use of the different metrics, applied to one example penetration. We generally begin by applying metrics that are most consistent and definitive for identification of borders and layers, then constrain possible solutions related to subsequent decisions where individual metrics used in isolation might fail. The particular metrics that are most reliable and which are prioritized differ depending on the particular boundaries that are being evaluated. However, as a general rule, CSD profiles were not necessary for layer identification and mostly were not used, with the exception of occasional cross-referencing of laminar response latencies. Finally, prior knowledge of the range of relative thicknesses and positions of layers based on anatomical studies (e.g. Figure 1A) is crucial for guiding the steps described below and can often be used to make very close estimates in cases where all metrics combined are still not sufficient to reveal a clear boundary. High resolution layer delineations were all manually derived for each of our penetrations following the guidance below. (See more examples in Figure S5, S6.)

**Figure 8.**
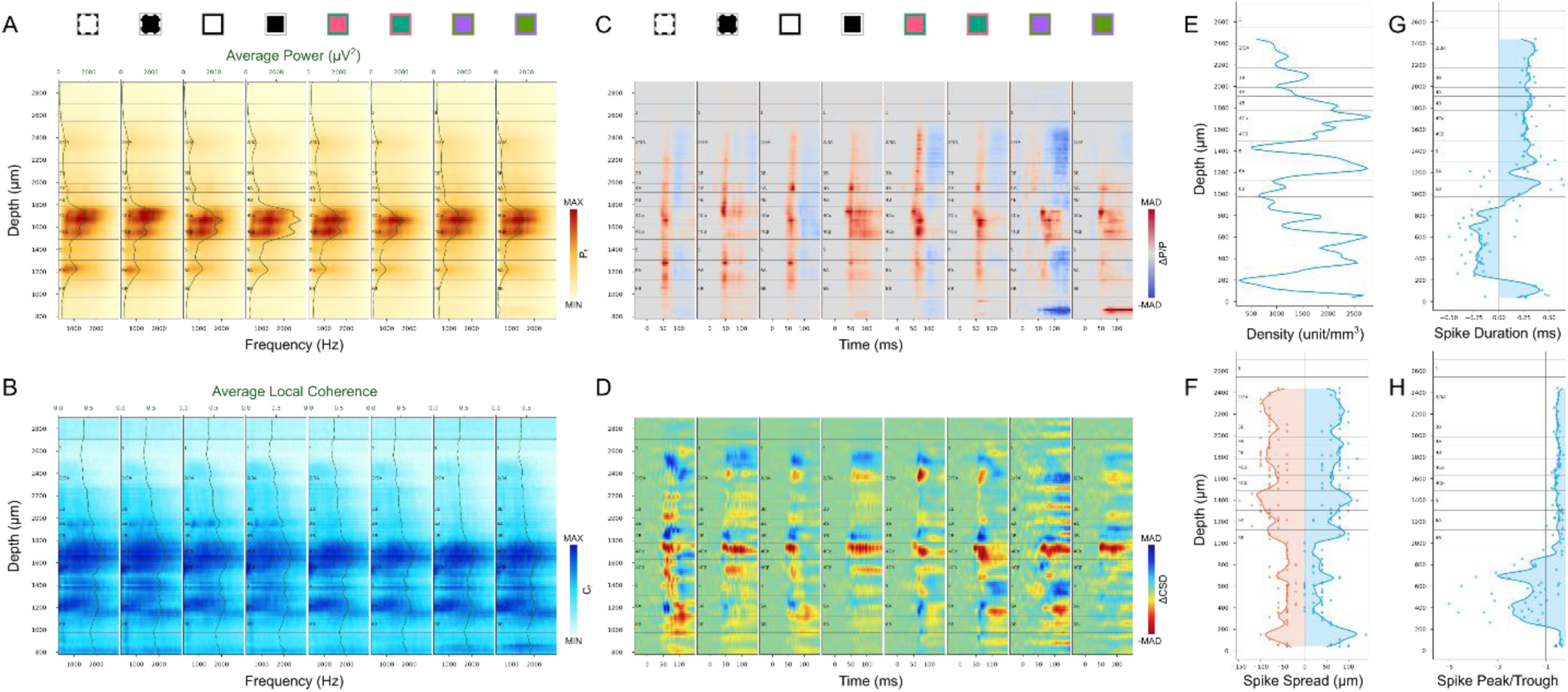
V1 layer delineation for an example penetration (A2) based on AP power spectrum (A), local coherence (B), instantaneous AP power changes (C), CSD changes (D), unit density (E), spike spread (F), spike duration (G) and spike peak-trough ratio (H). Conventions are the same as for corresponding plots illustrated in earlier figures.

### Top and bottom: L1/L2 and L6/WM

As a first step, it is very helpful to identify the top and bottom of the cortex. Given the known ranges of relative layer depths based on prior anatomical studies, knowledge of the total cortical depth and which electrode contacts are within the cortex provides an important constraint on subsequent possibilities. The L6/WM border can be reliably determined based on several measures that typically all corroborate one another. The most sharp and distinct transitions are the changes in spatial spread, duration, and peak-trough ratio of spikes (Figure 8F-H). This border can also be confirmed by the decreased coherence at the bottom of L6. Because neuron density increases steeply from L1 to L2, this border also has a clear transition that is most consistent across visual stimuli for relative power change (Figures 7, 8C), but is also clear for multiple stimulus conditions and across metrics based on AP power and coherence (Figures 5, 6, 8A, 8B). It is not unusual for unit density to fall to zero below the L1/L2 border (Figure 8E). This is likely because damage caused during initial electrode penetration through the pia can result in poor quality recordings such that units in L2/3A cannot be detected and isolated using spike sorting algorithms. But the other AP band measures (Power and Coherence) clearly show that the corresponding electrode contacts are surrounded by active neurons.

### Deep borders: L4C/L5, L5/L6 and L6A/L6B

The top and bottom of L5, corresponding to the L4C/L5 and L5/L6A borders respectively, are marked by both low anatomical cell density and an absence of LGN input to L5. Functionally, these changes are most consistently and easily recognized as sharp edges in AP power spectra, also corroborated by AP local coherence spectra (Figure 8A-B). L5 neurons also have the largest soma size in cortex that can be corroborated by the spike spread distribution (Figure 8F). The L6A/L6B border is identified from the decreases in AP power and local coherence that are found about halfway from the L5/L6A border to the L6/WM border. This relative width matches closely to the change in neuronal density that is used to define the L6A/L6B border in Nissl-stained anatomical specimens and also matches closely with the termination of magnocellular LGN afferents in L6B (*12*).

### Layer 4 sublayers: L4A, L4B, L4Cα, and L4Cβ

L4B, like L5, is a relatively cell sparse zone that lacks direct LGN input. Accordingly, the same criteria that distinguish the top and bottom borders of L5 can usually be used to identify the L4B borders. In particular, the L4B/4C border is almost always distinguished by a sharp decrease in AP power and local coherence (Figure 8A-B). Since L4B neurons are the projection targets of L4C neurons, their latencies are expected to be longer than that of L4C and sometimes this transition can be identified in ΔP/P profiles (Figure 7A, 8C). In the great majority of penetrations there is a clear thin band of signal corresponding to the expected location of L4A for at least one of the AP metrics under one of the stimulus conditions (e.g. Figure 8C). The top of this band is marked as the L3B/L4A border and the bottom as the L4A/L4B border. In 4 of 31 penetrations, no such electrical signature was seen to indicate the position of L4A. In these cases, L4B can be estimated to be half the thickness of L4C, and L4A can be estimated to be 50μm thick.

Anatomical studies reveal a clear change in neuronal density at the L4Cα/L4Cβ border and the thickness of L4Cα is invariably at least as large as for L4Cβ. The electrical signatures that best identify the L4Cα/L4Cβ border are due to the latency differences expected from the faster conduction velocity for magnocellular afferents to L4Cα versus parvocellular to L4Cβ. Between the metrics that reveal latencies, ΔP/P is more reliable and yields sharper transitions than ΔCSD profiles. When there is a clear latency transition in the middle of L4C (e.g. Figure 7A, 7B, 8C, 8D) this is usually marked as the L4Cα/L4Cβ border, but the position is always placed such that L4Cβ is not thicker than L4Cα because this is a consistent feature in anatomical observations. When there is no observable latency difference, the L4Cα/L4Cβ border is placed halfway between the top and bottom of L4C.

### Superficial borders: Pia/L1 and L2/3A/L3B

There are no clear electrical signals to distinguish the precise borders between the pia and L1 or between L2/3A versus L3B. Because L1 contains only a very low density of inhibitory neurons, it is often the case that few or no units are isolated above the L1/L2 border. Nevertheless, it is clear that no units can exist above the pial surface, so any neuron that is isolated above the L1/L2 border can be confidently assigned to L1. For simplicity, we simply assign the L1/pia border to a position ∼120-150µm above the L1/L2 border, as this is typical in postmortem histology and invariably incorporates any neurons that are recorded in L1.

The large span from the bottom of L1 to the top of 4A (L2/3) is relatively uniform in histological preparations and is often considered as a single layer. Nevertheless, there are clear differences in the connections and functional properties of neurons in the upper third of L2/3 versus the bottom third. For example, the CO staining intensity and the density of koniocellular S-ON afferent axons are both greater at the bottom of L2/3 (*12, 14*) and the local axonal projections of L4C neurons rarely extend above the bottom third of L2/3 (*22, 23*). While none of these differences relates to a distinct border, it is useful to identify neurons that are more superficial versus deep in L2/3, and we therefore simply mark the border between L2/3A and 3B at a position two-thirds of the distance between L1/L2 and the top of 4A.

#### Average metrics of V1 layer template

Figure 9 illustrates the laminar boundaries and thicknesses derived for all 31 electrode penetrations in our sample. Note that the overall thickness of the cortex in our physiological penetrations is somewhat more variable than is seen in postmortem histology, ranging from about 1.5 to 2.0mm. This is expected because there can be expansion due to brain swelling or shrinkage due to the placement of a coverslip onto the surface of the cortex in order to generate pressure and minimize brain movement; these influences can vary over the 4-5 days during which recordings were made from each animal. These influences are generally expected to generate proportional changes in laminar thickness across the cortical depth. An additional factor is that cortical thickness is known to decrease closer to the V1/V2 border and this is also seen in our data set (Figure S7A). Otherwise, there were no significant differences between monkeys or between ISI-identified functional domains (Figure S7B-D, Kruskal-Wallis Test, p>0.05). We used the complete set of 31 penetrations to make a proportional alignment and generate an “average” V1 layer template based on the average thickness of each layer (Figure 9B). This can be useful when laminar boundaries must be estimated in the absence of empirical data.

**Figure 9.**
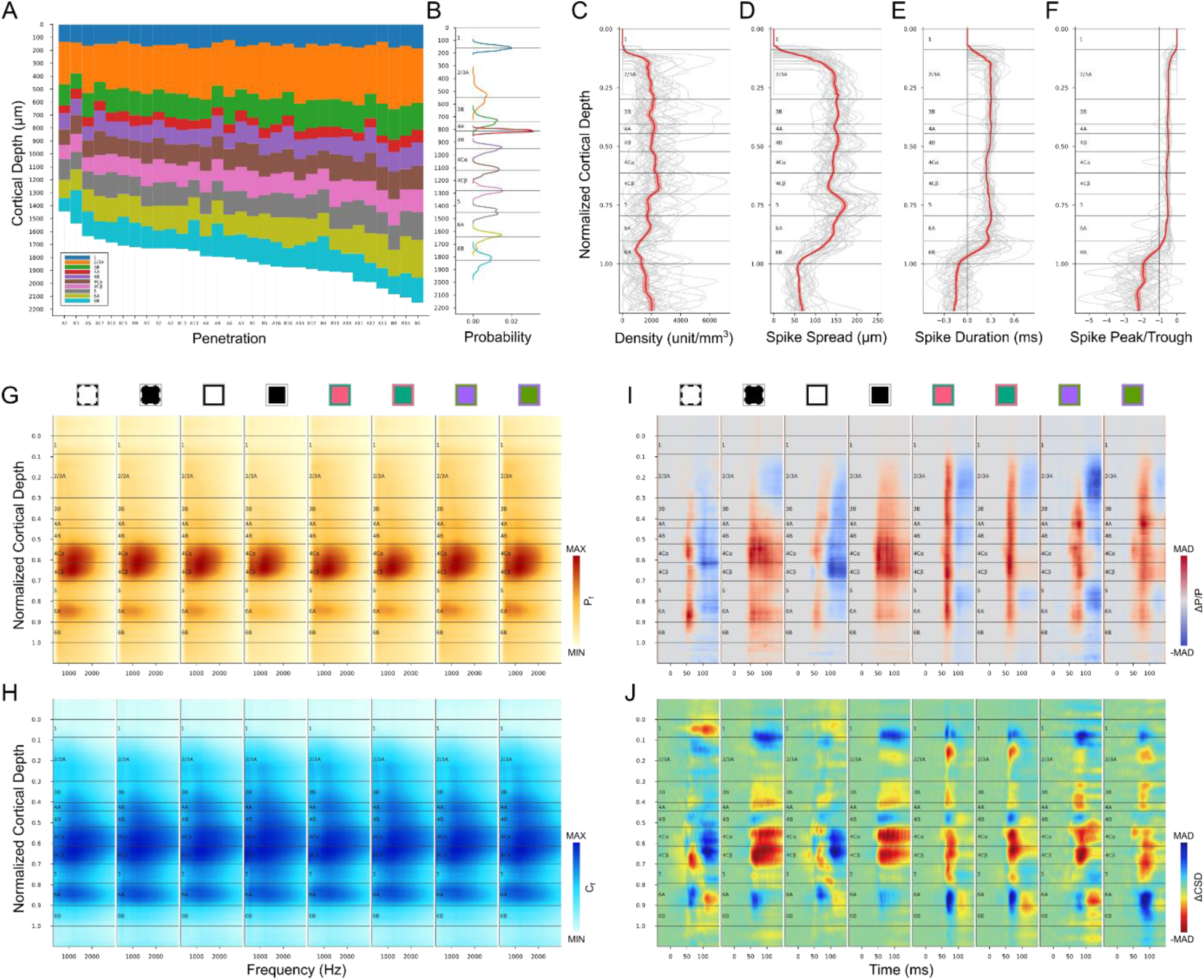
V1 layer template and average metrics across cortex. (**A**) Identified layers for 31 electrode penetrations aligned to the surface of cortex. (**B**) V1 layer template and probability density function for the thickness of each layer, aligned on the corresponding boundaries from the layer template. (**C-J**) Average depth distributions from 31 electrode penetrations. (**C-F**) First group of identification metrics (C: unit density, D: spike spread, E: spike duration, F: spike peak-trough ratio) on the normalized layer template (gray lines for each penetration, red line and shaded ribbon for Mean±SEM). (**G-J**) Second group of identification metrics (G: power spectrum, H: local coherence spectrum, I: ΔP/P, J: ΔCSD) on the normalized layer template. Conventions are the same as in previous figures.

We also generated averaged depth distributions for each metric (See Methods). The overall shape of the average unit density distribution (Figure 9C) matches that of the anatomical neuron density distribution reported in figure 6 of the study from Kelly and Hawken (*18*), with highest density in L4Cβ and decreased density in L5. Unit density also increases at the L5/L6 border, and in L4A compared to L4B and L3B. Due to the limited sampling volume of a linear probe, tissue damage, and spike sorting capabilities, the unit densities are far less than neuron densities calculated from immunohistology, and density contrast across layers is greatly reduced. The average spike spread (Figure 9D) matches the histological observation that L5 neurons have the largest cell bodies while L4Cβ neurons are the smallest. Note that the decreased spike spread in L4A is also expected from the small cell bodies in that layer. The location of the white matter boundary has a marked drop in spike spread corresponding to the transition from spikes recorded from cell bodies versus axons. This transition can also be seen in average distributions of spike duration (Figure 9E) and spike peak-trough ratio (Figure 9F).

The average power spectrum (Figure 9G) and local coherence (Figure 9H) distributions clearly show the expected decrease in L5 relative to L4C and L6A. The pattern was also stable across different stimulus conditions implying closer relation to anatomical properties than functional ones. The average relative power change (Figure 9I) shows strongest activation in L4A using S/(L+M) cone-opponent colors. It also can be seen that the latencies of 4Cα responses were shorter than for 4Cβ. Although CSD was rarely used in our layer identification process, the average ΔCSD profiles (Figure 9J) were not far from the expectations based on many previous results. It seems that switching from white to a black screen evoked the strongest increase of current sinks in L4C. For all stimuli, L5 shows increased current sinks while L6A shows increased current sources. L4B shows increased current sources, while L4A/L3B shows increased current sinks.

## Discussion

### Laminar identification in primate V1

We have described a set of metrics based on high-resolution AP-band (0.3 – 10kHz) signals from Neuropixels probes that can be used to identify cortical layers in primate V1 at higher resolution and with greater accuracy than using LFP band signals (0.5 – 500Hz) and CSD analysis. Our methods capitalize on properties of electrical signals that can extract three key features of V1 laminar organization: 1) anatomical differences in the size and density of cell bodies; 2) functional differences in the LGN afferents and recipient neurons in different layers; 3) laminar termination patterns of functionally distinct LGN afferents. Accordingly, anatomical differences can often be extracted even without the use of visual stimuli, while functional differences require activation with different visual stimulus conditions and often also incorporate consideration of response latencies.

### Limitations of LFP signal and CSD analysis

Here, we have shown that very close (20μm) electrode spacing and increased electrode density does not necessarily result in higher resolution identification of laminar boundaries based on CSD profiles. Nevertheless, low density electrode arrays and CSD methods have been used for cortical laminar identification in various species and cortical areas (*24, 25*), in both awake (*8*) and anesthetized recordings (*9*). Although the use of CSDs has evolved from simple second spatial derivative to model based inverse methods (*7, 26, 27*), the underlying assumptions of the forward problem remain the same: continuous CSD distribution, quasistatic electrodynamics and ohmic extracellular media (*6*). When electrode spacing is close to or less than 20μm, the assumption of continuous and macroscopic CSD distribution might not be valid, and models of CSD distribution based on discrete microscopic sinks and sources may be more appropriate. The alternating sink and source patterns of CSD profiles (Figure 2A) derived from 20µm spacing LFPs may reflect the microscopic CSD dipoles of neurons. Thus, an appropriate spatial averaging or down sampling of electrodes was needed to produce smooth, useful CSD profiles in which regions of sinks and sources can be potentially related to cortical layers or sublayers. The sizes of smoothed sink or source regions are typically most suitable for matching the thickness of layers and sublayers when electrode spacing is around 50μm. This implies that further reductions in probe spacing will not improve the ability of CSDs to identify cortical layers.

The CSD profile represents spatial and temporal patterns of electrical activities across layers, thus it naturally depends on the structure and dynamics of functional cortical circuits. We have shown that for any given electrode penetration, differential activation of geniculocortical pathways using different colors or achromatic stimuli generates distinct CSD profiles. For example, CSD profiles generated in response to S-cone stimuli can have relatively large sinks (depolarization) in layers 4A and 3B, similar to those generated by directly activating koniocellular LGN neurons using electrical and optogenetic stimulation (*9*). CSD profiles generated from black and white stimuli also show dramatic differences. Flipping the screen from white to black consistently generated strong early sinks that roughly matched layer 4C. On the contrary, flipping from black to white sometimes generated early sources (hyperpolarization) instead of sinks (Figure S2D) making it less reliable than a black stimulus. The large white stimulus has been reported to induce sustained inhibition which was absent in black stimulation (*20*). This suggests that CSDs from black and white stimuli should be analyzed separately for layer identification.

To increase spatial resolution of LFP signals, prior studies have restricted LFP analysis to the gamma band (30-100Hz), which should theoretically spread over shorter distances than when lower frequencies are also included (*17*). Indeed, when this is used to identify monkey V1 layers, a layer 4C sink is detected with a somewhat sharper cutoff at the likely L4C/L5 border than for the CSD (*10*). When we apply this analysis to our data collected with Neuropixels probes (Figure S8A, C), we see similar patterns to those previously reported (*10*); however, the ability to accurately detect laminar boundaries or to definitively identify the precise borders of layer 4C are not comparable to what can be achieved using AP band signals from high-density Neuropixels probes.

LFP gamma band coherence has also been used (*10, 24, 28*) to categorize ensembles of neuron populations, and several of them have been shown to be confined to specific layers. We analyzed local coherence using gamma LFP (Figure S8B, D) and find that the ability to detect laminar boundaries is also much poorer than signals based on AP band coherence, even when using high density Neuropixels probes. It is unclear what mechanisms generate the laminar coherence in gamma LFPs, however, our clearer laminar signals based on AP band local coherence indicates that this is likely due to the sharper falloff of high frequency signals, which can be exploited using only the densely spaced Neuropixels electrode contacts.

### AP signals have high spatial resolution

In contrast to the inherent low resolution of LFP-based signals, signals based on AP band metrics have the potential to allow higher resolution laminar detection when using high-density probes. The various AP-band metrics that we have extracted and used to allow laminar identification include: unit density, spike spread, duration and peak to trough ratio, power spectrum, local coherence spectrum and temporal power change profile. Most of these measures could be analyzed based on spontaneous activity and therefore might be utilized in brain areas other than V1, such as higher cortical areas where the additional signals derived based on visual response latencies or laminar differences in functional input properties are not as prevalent. Indeed, in rat hippocampus, it has been demonstrated that high frequency power (≥300Hz) can be used to differentiate different layer structures (*28*).

We have described the hypothetical generative models of AP band power and local coherence based on superposition of individual spikes. In the case of similar order of firing rate across layers, the depth distributions of AP band power and local coherence mainly represent depth distributions of neuron density which is the most definitive anatomical property for layer delineation. Therefore, AP band power and local coherence should be much less dependent on stimulus, as demonstrated in our results. Moreover, they remained largely unchanged even in the baseline period (Figure S9) in which cortical circuits were believed to have been stabilized following stimulus-induced activity. Since AP power and local coherence were homogenous across frequencies, they could also be first averaged across frequency, then shown for consecutive trials (Figure S10). This opens the possibility of online layer estimation without visual stimulation.

The improvements in layer identification that we have demonstrated using high-resolution AP-band signals and high-density probes suggest that still greater improvements might be possible. The estimation of neuron density profile, for example, should be more accurate when more neurons can be detected and isolated by spike sorting. This can be achieved by decreasing electrode spacing or by using multiple shanks. Neuropixels 2.0 probes (*29*) have even closer electrode spacing at 15μm and up to four shanks; thus, the spike-sorted units within and across layers should allow even more accurate estimation of neuron density distribution and better laminar delineation. Furthermore, new development of spike sorting and drift correction algorithms (*30*) that incorporate detailed modeling of spike spread could better estimate the three dimensional positions of units, leading to better unit identification and drift estimation, and may also estimate soma properties, giving additional anatomical information useful for laminar differentiation.

In addition to stimulus independent measures, we have also incorporated responses to visual stimuli for our laminar analyses based on power spectra, local coherence spectrum and stimulus-evoked temporal changes in AP power. We have demonstrated the ability to detect functional distinctions that are peculiar to monkey V1. For example, we found that the most effective way of revealing the thin layer 4A was to use colors that modulate only LGN S Cone inputs. The response latency differences between 4Cα and 4Cβ are also useful to separate these two otherwise ambiguous layers. On the other hand, the L-M cone opponent colors were expected to much more strongly activate layer 4Cβ, however, substantial activation with little delay was observed across supragranular, granular and infragranular layers, implying complex intralaminar processing of color in V1 (*19*).

### Consideration of electrode drift

There are several factors that can cause the relationships between electrode contacts and laminar borders to change over time. These include both changes on a fast time scale (such as head movement, heart rate, or breathing) and slower drift, such as expected from brain swelling. In addition, there can be either “rigid” uniform movements that simply shift the locations of contacts relative to the tissue, or non-uniform changes that might differentially affect the relationships at different locations on the probe (such as compression of superficial layers during initial introduction of an electrode).

In our experiments, electrode movement was minimized and often not detected, resulting in stable recordings (Figure S12A). This is primarily because we used light compression of the tissue by placement of a coverslip at the surface, which largely eliminates fast movements from heartbeat and breathing. Furthermore, the animals are anesthetized and paralyzed, preventing changes due to head movements. Nevertheless, slow drift was detected in some of our recordings, likely due to swelling of brain tissue over time (Figure S12B). We corrected for drift using Kilosort (*31*) version 3 on concatenated AP data produced by CatGT (See Methods). This approach is based on signals in the AP band which, as described above, are expected to have better spatial resolution than LFP signals.

In contrast, prior studies have used LFPs to track probe motion and in turn correct for motion in human cortex (*32*). However, LFPs may not be a stable or accurate indicator of electrode drift because the depth pattern of LFP might be changed by external stimuli or internal spontaneous activities, dissociating the temporal change of LFP patterns from probe motion. On the other hand, spike sorting algorithms (e.g. Kilosort 3.0) should be able to use spike amplitudes to track tissue displacement relative to probe contacts with higher accuracy than LFP. While this requires sampling sufficient numbers of spikes, introducing tradeoffs between spatial resolution and sensitivity, these tradeoffs are minimized by the use of high-density electrode arrays, such as Neuropixels.

Regardless of whether laminar boundaries are estimated from CSDs, LFPs, or AP signals, matching unit positions to laminar locations should use motion correction for both rigid and non-rigid sources. We found that strict reliance on a rigid model for motion correction with Kilosort version 3.0 failed to accurately identify slow drifts in some of our penetrations (Figure S12E). We therefore also introduced additional methods to allow for non-rigid drift (See Methods).

### Towards a universal in vivo pipeline for cortical layer identification

In this study, we developed AP metrics that can be used to estimate several important anatomical and functional properties of cortical layers. To accurately identify each layer, multiple metrics need to be combined, since no single metric is sufficient to reveal all layers. Here, layer boundaries were manually drawn, but automatic/semi-automatic algorithms can be constructed. Template matching algorithms have been developed to automatically align probe electrodes to cortical layers (*33*). Since multiple metrics would need to be incorporated, complicated regression algorithms would potentially be needed to satisfy multiple objectives, and the layer thickness template we derived (Figure 9B) could be used as a prior to penalize large deviations from the template. On the other hand, supervised and unsupervised machine learning algorithms might be able to fully exploit all metrics and provide less biased estimates (*34*). However, large datasets and annotations may be needed to train and test the artificial neural network models to ensure that they can operate at an accuracy level no less than human operators.

The laminar organization of cortical circuits is thought to be related to functional circuit mechanisms underlying many cognitive functions. Thus, assignment of in vivo recorded neurons to cortical layers could facilitate anatomical and functional analysis of information processing in cortical circuits in areas beyond the monkey primary visual cortex. The expectation that evoked initial sinks correspond to input layer 4 is likely limited to primary sensory areas where the main feedforward input from the thalamus terminates densely in that layer and can be driven robustly by a single stimulus. It has been shown that the location and thickness of initial sinks could vary across multiple mouse visual cortices, making it hard to identify layer 4 according to initial sinks (*35*). Moreover, in cortical areas that do not directly process sensory information, such as prefrontal cortex, the input layer 4 may not be easily activated by external stimuli, rendering CSD methods unusable. The AP metrics estimating mainly anatomical properties could overcome the structural and functional variations across different cortical areas and species, leading to a general solution for in vivo layer delineation. Future simulation studies based on biophysically plausible cortex models would not only reveal the applicable conditions and limitations of the AP metrics we described, but could also lead to identification of new metrics to more accurately estimate essential anatomical properties, such as neuron density and soma size/shape, that have been accepted for universal delineation of cortical layers.

## Materials and Methods

### Animal Surgery

Two rhesus monkeys (*M. mulatta*, adult, male) were used for the acute experiments. All procedures using live animals were approved by the Institutional Animal Care and Use Committee (IACUC) at The Salk Institute for Biological Studies. Detailed surgical procedures have been described previously(*36*).

Animals were injected intramuscularly with atropine sulfate (0.02 mg/kg), and then anesthetized with ketamine (10 mg/kg). During surgery, animals were kept under anesthesia (1-2.5% isoflurane in O_2_) and sterile conditions. Tracheostomy was performed, and animals were mechanically ventilated. EKG, SpO_2_, end tidal CO_2_, and body temperature were all monitored throughout the experiments. Lactated Ringer’s with 5% dextrose was given intravenously. Antibiotics (Cefazolin, 25 mg/kg, i.v.) were administrated every 12 hours. Mannitol (20%, i.v., 200mg/kg) was given slowly 30 minutes before craniotomy was performed. Dexamethasone (1-2 mg/kg, i.v.) was given daily.

A large craniotomy (∼25mm diam.) and durotomy were performed over the primary visual cortex, and a customized recording chamber was cemented onto the skull. A coverslip was mounted to reduce brain movement. Two silver wires were implanted from which EEG signals were monitored to assess anesthetic depth. Anesthesia was gradually transitioned from isoflurane to sufentanil citrate (2.0-8.0 µg/kg/hr, i.v.), and vecuronium bromide (0.05-0.2 mg/kg/hr i.v.) was administered to paralyze the animal after the chamber was implanted. Sufentanil dosage was adjusted to ensure slow wave EEG.

### Extracellular Electrophysiology

A Neuropixels 1.0 probe (IMEC, Leuven) was held by an oil hydraulic manipulator (Narishige International USA, Inc.), which was attached to the chamber. The probe was aligned perpendicular to cortex and slowly inserted around 3mm below brain surface through a small hole (∼1mm diameter) in the coverslip. Then, the hole was covered with saline to prevent drying. Reference and Ground of the probe were soldered together and connected to the chamber. Extracellular electrical potentials from 384 channels were sampled at 30kHz and streamed to a computer through the Neuropixels acquisition module (IMEC, Leuven) within a PXIe system (PXIe-1071 chassis, PCIe-8381, PXIe-8381, PXIe-6341, National Instruments, Corp.). Data acquisition was controlled by SpikeGLX (https://billkarsh.github.io/SpikeGLX). Raw data were filtered and saved separately to an action potential band (AP band, 0.3 – 10kHz, 30kHz) and a local field potential band (LF band, 0.5 – 500Hz, 2.5kHz). Recordings started around 30 minutes after probe insertion when responsive visual fields were manually mapped. The time interval between different penetrations of the same animal was at least 3 hours.

### Visual Stimulus Presentation

Visual stimuli were generated and presented using Experica (https://experica.org) on an LCD monitor (ROG SWIFT PG279Q, Asustek Computer Inc.) with a refresh rate of 144Hz. The monitor was positioned 57cm in front of the animal’s eyes, which were dilated with 1% atropine sulfate and 0.5% proparacaine hydrochloride ophthalmic solution. Both eyes were fitted with contact lenses to focus on the screen.

Prior to each experiment, the luminance of the red, green and blue primary colors of the monitor were measured using a spectroradiometer (PR-701, Photo Research, Syracuse, NY) and linearized by a color lookup table. Then, the spectra of red, green, blue primary colors were measured at resolution of 2nm. Three color pairs on the three axes of DKL color space with maximum achievable cone contrast: L+M luminance (white/black), L/M cone opponency (red/green), and S/(L+M) cone opponency (purple/lime), were calculated (https://github.com/Experica/ColorLab.jl) based on the spectra of red, green, blue primary colors and the 10° cone fundamentals (*37*). The two colors of a pair were presented alternatively, each for 1.5 second on full screen and repeated 100 times. All the color pairs were presented to dominant eye, whereas only the L+M luminance color pair was presented to non-dominant eye. The total time of presenting all stimuli described here was in 30 minutes or less.

### Data Analysis

Data were first processed (https://github.com/Experica/NeuroAnalysis) in MATLAB (Mathworks Inc., MA), then analyzed (https://github.com/Experica/NeuroAnalysis.jl) in Julia (https://julialang.org).

In order to correct probe drift during whole recording session for each electrode penetration and assign consistent ID for each unit in spike sorting, all AP band data of one penetration were concatenated in time order by CatGT (https://billkarsh.github.io/SpikeGLX). In the process, a subsampling method was used to demultiplex (TShift option) channels that shared the same ADC of the probe, then a 300Hz high pass Butterworth filter of order 3 was applied to remove offset, and finally a global common average referencing was used to remove shared noise.

### Drift correction and spike sorting

Kilosort (*31*) version 3 (https://github.com/MouseLand/Kilosort) was used on concatenated AP data produced by CatGT, and the sorting results were curated in Phy (https://github.com/cortex-lab/phy). Prior to spike sorting, whitened data were saved to a separate file and a rigid registration method was used to correct probe drifting in the file. In the anesthetized state, most of our penetrations showed little to no drift.

For a few penetrations in which significant drift occurred, non-rigid drifting was found prominent, and a non-rigid registration method (imregdemons in MATLAB) was used instead of the rigid one in Kilosort if the rigid drift correction result was not satisfactory (Figure S12). The putative double counted spikes (inter-spike-interval ≤ 0.35ms) within a cluster and across nearby (within circle of 60µm radius) clusters were removed before manual curation in Phy. Clusters with low mean firing rate (≤ 0.5 spike/second) were regarded as noise. On average, around 200 single-unit or multi-units can be resolved for one penetration.

### Spike waveform feature and unit density

The spike waveforms of a unit were calculated by averaging randomly chosen spike AP segments (<1000) of the unit. The heights (peak subtracts trough) of spike waveform on different channels were used as weights to estimate (center of mass) unit position. The channels used in weighted average of channel positions were chosen within a circle of 65µm radius, centered on the channel of the largest spike height. The upward and downward spread of spikes were defined as the largest vertical distances between the channel with the maximum spike height and the closest channels with spike height less than 15% of the maximum spike height (*38*). Using the spike waveform of maximum height, the spike duration was defined as the time of peak subtracting time of trough, and spike peak-trough ratio was defined as the ratio of peak and trough amplitudes.

The unit density was estimated by assuming a 100µm radius cylinder as the unit detection volume of the probe. Depth distribution of unit density was derived by counting number of units in a 40µm segment of the cylinder moving along the probe with 20µm step and then smoothed by a gaussian filter (σ=28µm). A single unit was counted as one neuron, whereas a multi-unit was counted as 1.2 neurons. This value was empirically chosen based on a general impression of contamination rate of multi-units and likelihood of sharing the same units among adjacent multi-units. Since the percentage of multi-units was low (∼20%) in most of our penetrations, varying this value from 1 to 2 did not substantially change the signature shape of unit density distribution. The same methods of moving window and smoothing in unity density estimation were also used to estimate depth distributions of upward spread, downward spread, spike duration and spike peak-trough ratio.

### Current source density

The analysis time window of each stimulus presentation was chosen as [-40, 150] millisecond around stimulus onset time 0, where [-40, 10] was used as baseline window and [10, 150] as response window. These definitions also apply for the power spectrum, local coherence spectrum and temporal power change described subsequently. The local field potentials in analysis window were first notch filtered to remove electrical line noise (60Hz), then bandpass filtered in 1-100Hz and down sampled to 1kHz. LFPs with the same depth were first averaged while excluding the reference channel, then trial averaged to produce a series of LFP traces across cortex with a depth resolution of 20µm. Current source densities were calculated for each stimulus from the series of LFP traces. Specifically, the inverse CSD method (*7*) was used, and the

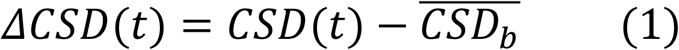

current source densities were modeled as infinitely thin disc with 500µm radius. The stimulus induced CSD changes (ΔCSD) were defined as CSD subtracting the temporal average CSD in the baseline time window.

### AP band power and coherence spectrum

In order to keep aligned with positions of sorted units, analysis was based on the file containing whitened, drift-corrected AP band data produced by Kilosort before spike sorting. The AP band data were unwhitened, bandpass filtered in 300-3000Hz, and down sampled in 10kHz. Power spectral densities across frequencies (*P*_*f*_) in the response window ([10, 150]ms) were estimated by multi-taper method. The instantaneous

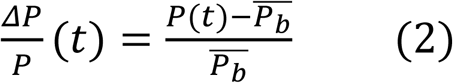

powers in analysis window ([-40, 150]ms) were evaluated by mean of squares (RMS^2^) in a 10ms sliding window. In the same way as for LFP, power spectral densities and instantaneous powers were averaged across same depth and trials. Depth distribution of power spectrum was smoothed by gaussian filtering (σ=20µm). The stimulus induced instantaneous power changes (ΔP/P) were defined as relative change to average instantaneous power in the baseline window ([-40, 10]ms).

Coherence is defined as follows:

where S_ii_(f) and S_jj_(f) were power spectral densities of channel i and j respectively, and S_ij_(f) was cross power spectral density of channel i and j. The local coherence spectrum (*C*_*f*_) for each channel was evaluated

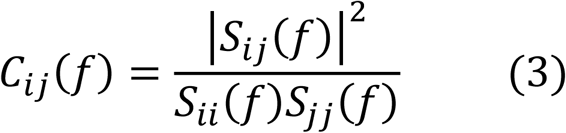

using weighted averaging where pairing channels were confined within a circle of 55µm radius, and distance between pairing channels was used to obtain gaussian weight (σ=25µm). Coherences in the response window ([10, 150]ms) were calculated by multi-taper method and then averaged across same depth and trials as described for LFPs to produce the distribution of local coherence spectrum along the probe.

### Layer normalization template

Several penetrations were excluded from the dataset when: (1) the health of the cortex had deteriorated, usually due to prolonged duration of maintenance under anesthesia (A17, B18); (2) the penetration was very close to the V1/V2 border, where the whole thickness of cortex is thinner than other regions of V1 (A7); (3) an error was made in assignment of the dominant eye, resulting in lack of data under several stimulus conditions (A8). The average thickness of each layer was used to build the V1 layer template, then the thickness of each layer for each penetration was linearly scaled to match the thickness of the corresponding layer in the V1 layer template. After mapping of depth coordinates of each penetration to the layer template, the depth distributions of identification metrics were interpolated (Cubic Spline) in depth coordinates of the template so that the average metrics on aligned layer template can be calculated. For power spectrum, local coherence spectrum, instantaneous power changes, and ΔCSD, profiles of each stimulus presentation for each penetration were normalized before averaging across penetrations.

## Acknowledgements

We thank Anupam Garg for discussions and suggestions for Neuropixels setup and data analysis, Bill Karsh for upgrading SpikeGLX to accommodate our experimental environment. We also thank Angel Macias and Angelo Salinda for their assistance in surgery and recording.

## Funding

NIH grants EY022577 (E.M.C)

NS105129 (E.M.C.)

Pioneer Fund (P.L.)

The Fundamental Research Funds for the Central Universities, No. 226-2024-00022, 2023ZFJH01-01, 2024ZFJH01-01 (P.L.).

The Non-profit Central Research Institute Fund of Chinese Academy of Medical Sciences, No. 2023-PT310-01 (P.L.).

## Author contributions

Conceptualization: LZ, PL, EC

Data Collection: LZ, PL

Methodology: LZ

Writing: LZ, PL, EC

## Data availability

All data necessary to support the paper’s conclusions are present in the main text, supplemental information, or available from the corresponding author upon reasonable request. The results presented here are based on more than 5TB of raw data and include data measuring activity during stimulus conditions that are not relevant to the present manuscript. The authors are presently conducting further analyses of that data which will be published at a later time. These data are not structured to allow data relevant to this manuscript to be separated from that which is reserved for future publication.

## Code availability

Customized code used to perform the analyses can be found at https://github.com/babaq/EPhys_V1_Layer and https://github.com/Experica.

---

**Figure S1.**
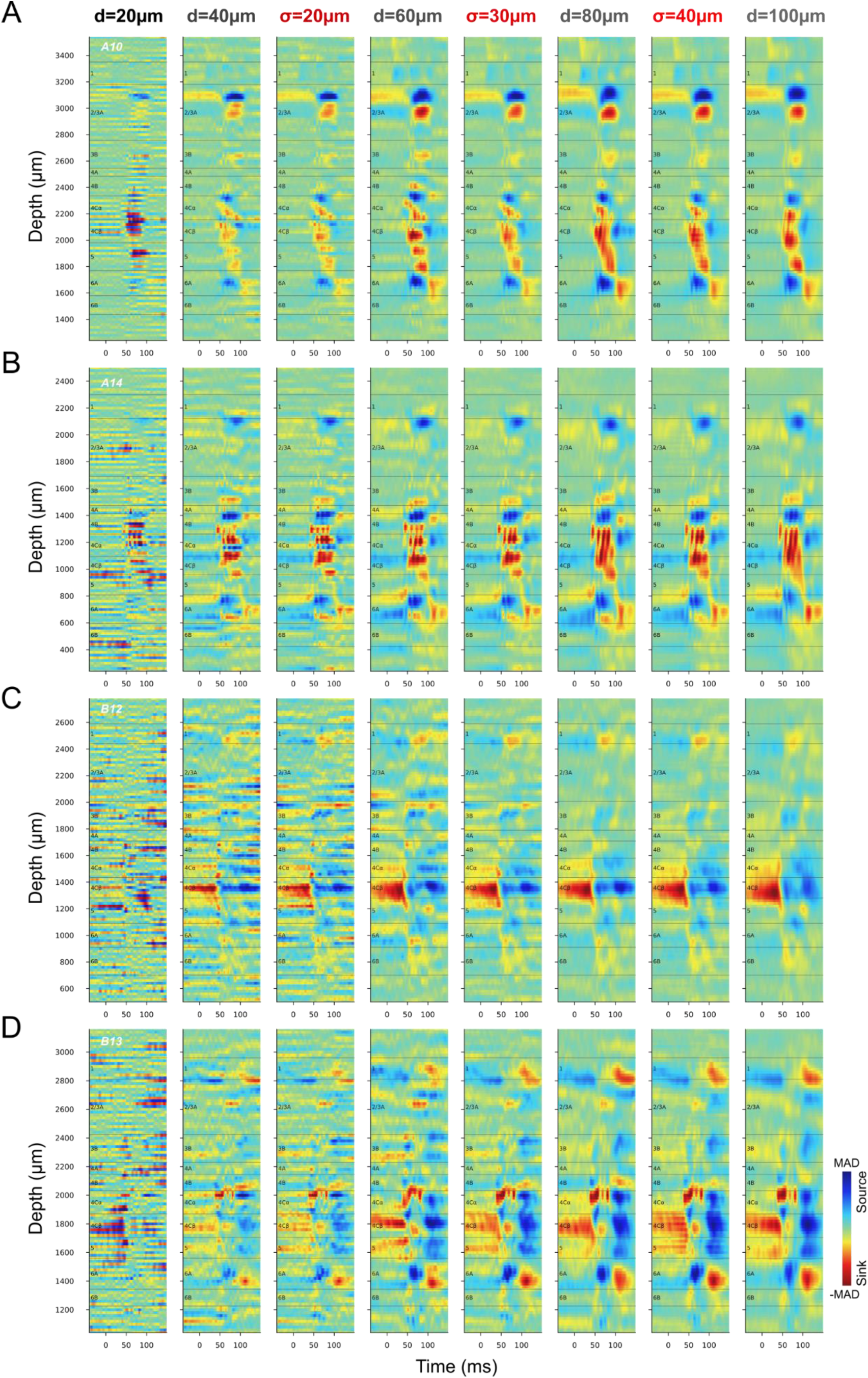
Comparing spatial averaging to downsampling for CSD profiles evoked by flipping the black screen to the white screen through the dominant eye. The same four penetrations in Figure 2E-H are shown here (A: A10, B: A14, C: B12, D: B13). In the downsampling process, LFPs from electrodes with increasing vertical spacing (d=20, 40, 60, 80, 100µm, gray) were used to calculate CSDs, then they were interpolated (Cubic Spline) to the vertical resolution of 20µm. In the spatial averaging process, CSD profiles of 20µm spacing were Gaussian filtered with increasing width (σ=20, 30, 40µm, red). The CSD profiles were independently color-mapped according to the maximum absolute deviation (MAD) in each profile.

**Figure S2.**
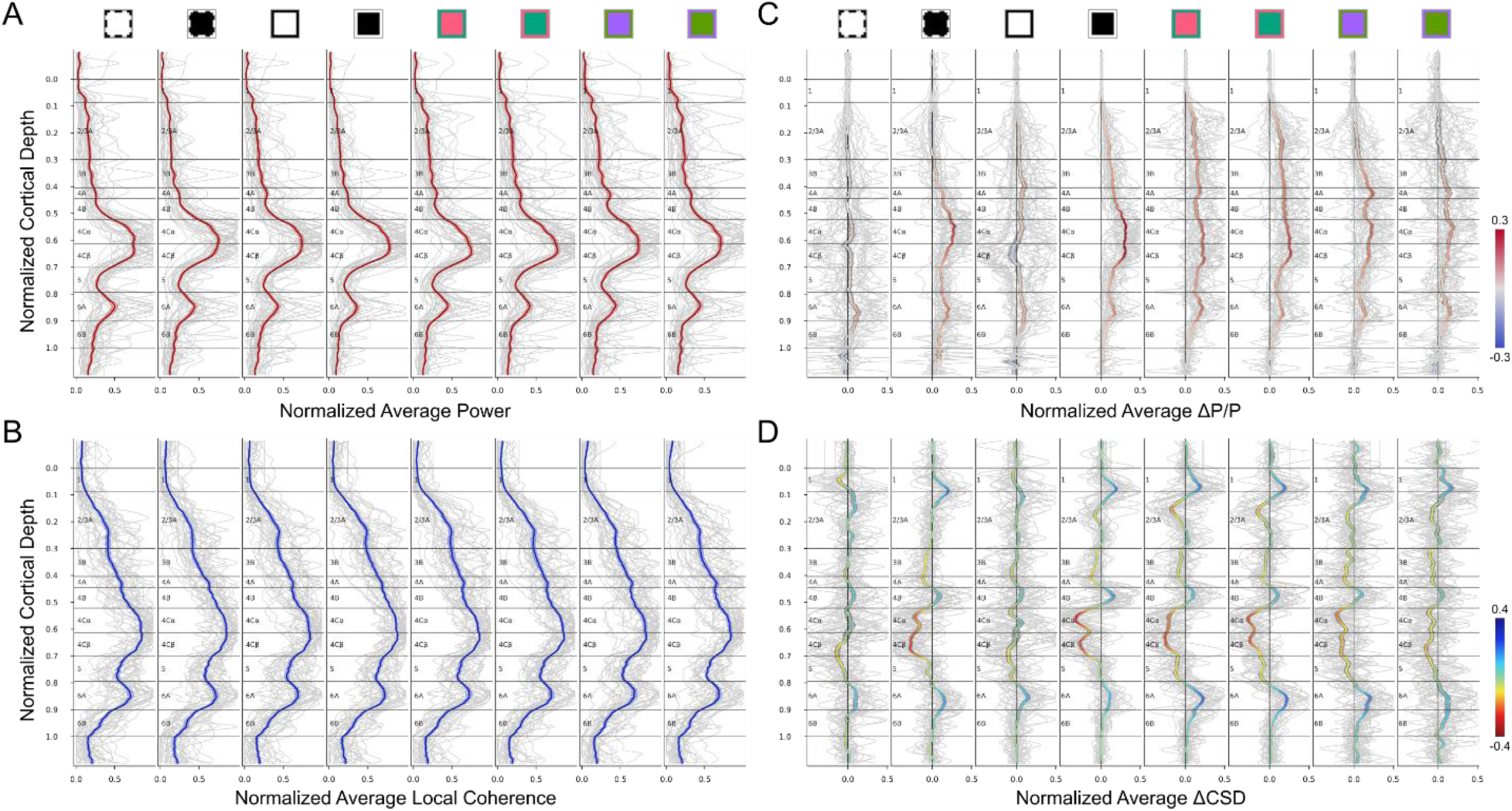
Average Metrics in V1 template. (**A**) Normalized average power for each stimulus. Gray lines are for each penetration, and red lines and shaded ribbons show Mean±SEM. (**B**) Normalized average local coherence for each stimulus is shown similarly to (A). (**C**) Normalized average ΔP/P in time window: [30, 100]ms after stimulus onset. (**D**) Normalized average ΔCSD in the same time window of C. The color-mapped lines and shaded gray ribbons in C and D show Mean±SEM. Square Markers are the same as in the figures of this article.

**Figure S3.**
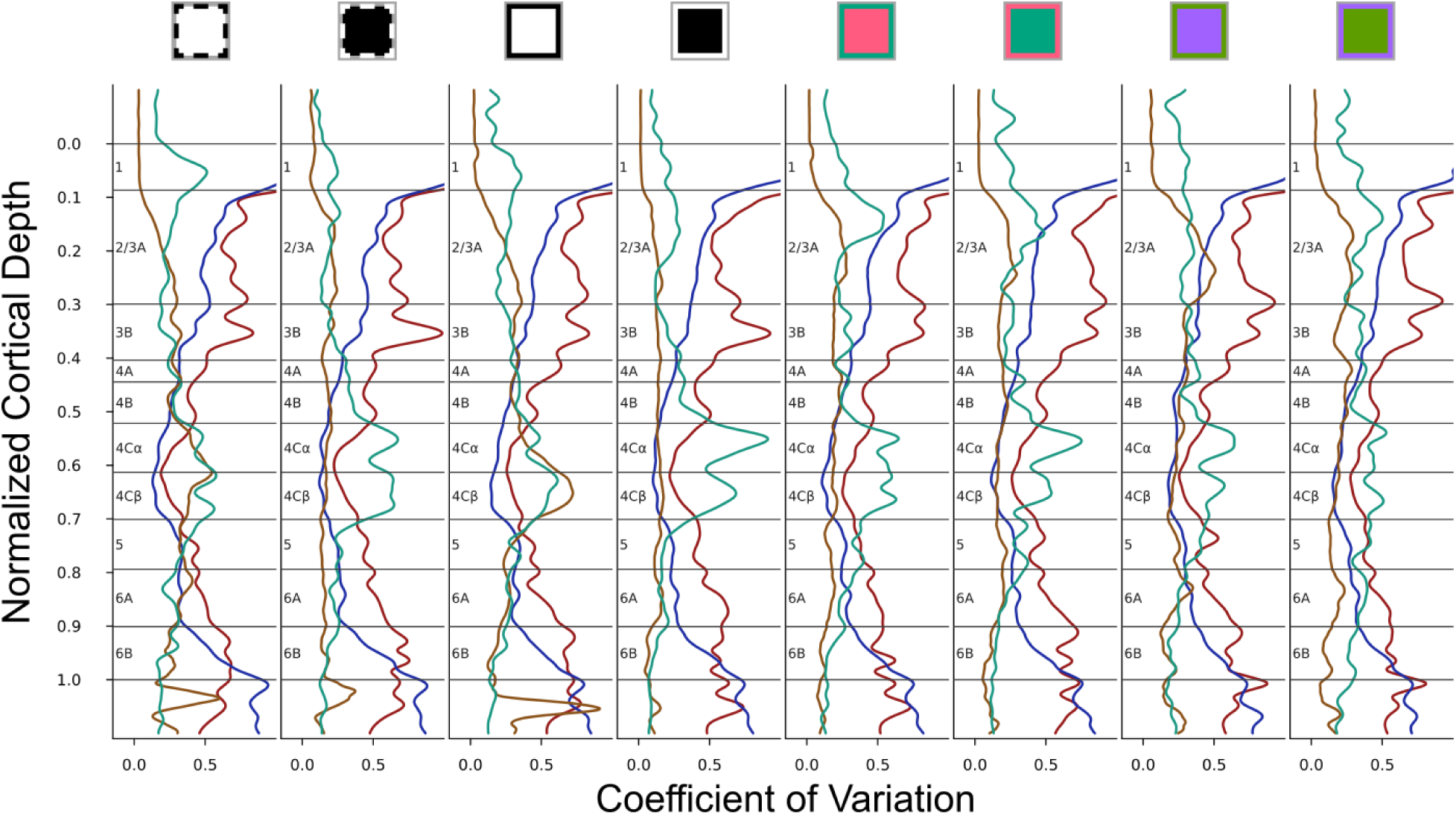
Coefficient of Variation. Variability of normalized depth profiles (gray lines in Figure S2) across penetrations is accessed by coefficient of variation (STD/Mean) for power (red), local coherence (blue), ΔP/P (orange), and ΔCSD (green) in response to different stimulus conditions. Square Markers are the same as in the figures of this article.

**Figure S4.**
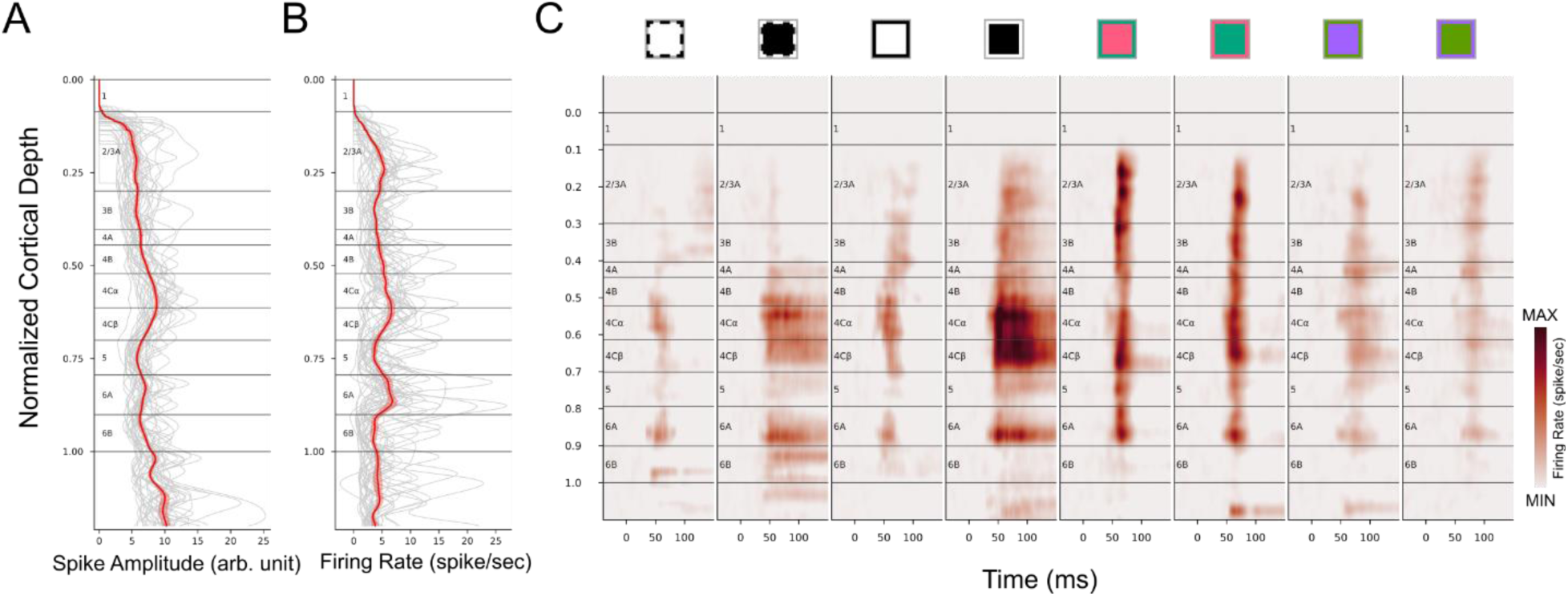
Depth profile of spike amplitude and firing rate. (**A**) Spike amplitude. (**B**) Firing rate during the whole recording session of a penetration. The stimulus set includes static/drifting, achromatic/chromatic gratings. (**A-B**) gray lines for each penetration, red line and shaded ribbon for Mean±SEM. (**C**) PSTH on the normalized layer template. Square Markers are the same as in the figures of this article.

**Figure S5.**
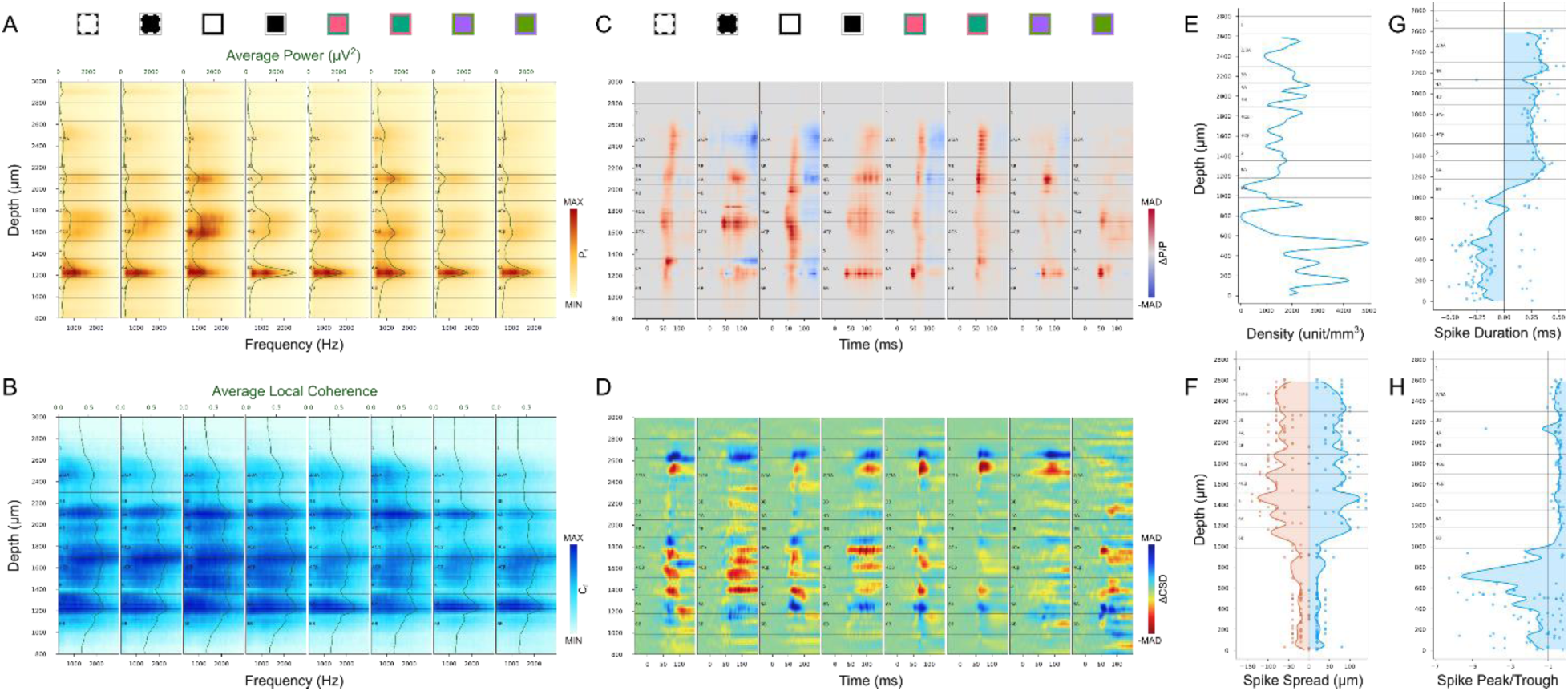
V1 layer delineation for example penetration (A3) based on power spectrum (A), local coherence (B), instantiations power changes (C), CSD changes (D), unit density (E), spike spread (F), spike duration (G) and spike peak-trough ratio (H).

**Figure S6.**
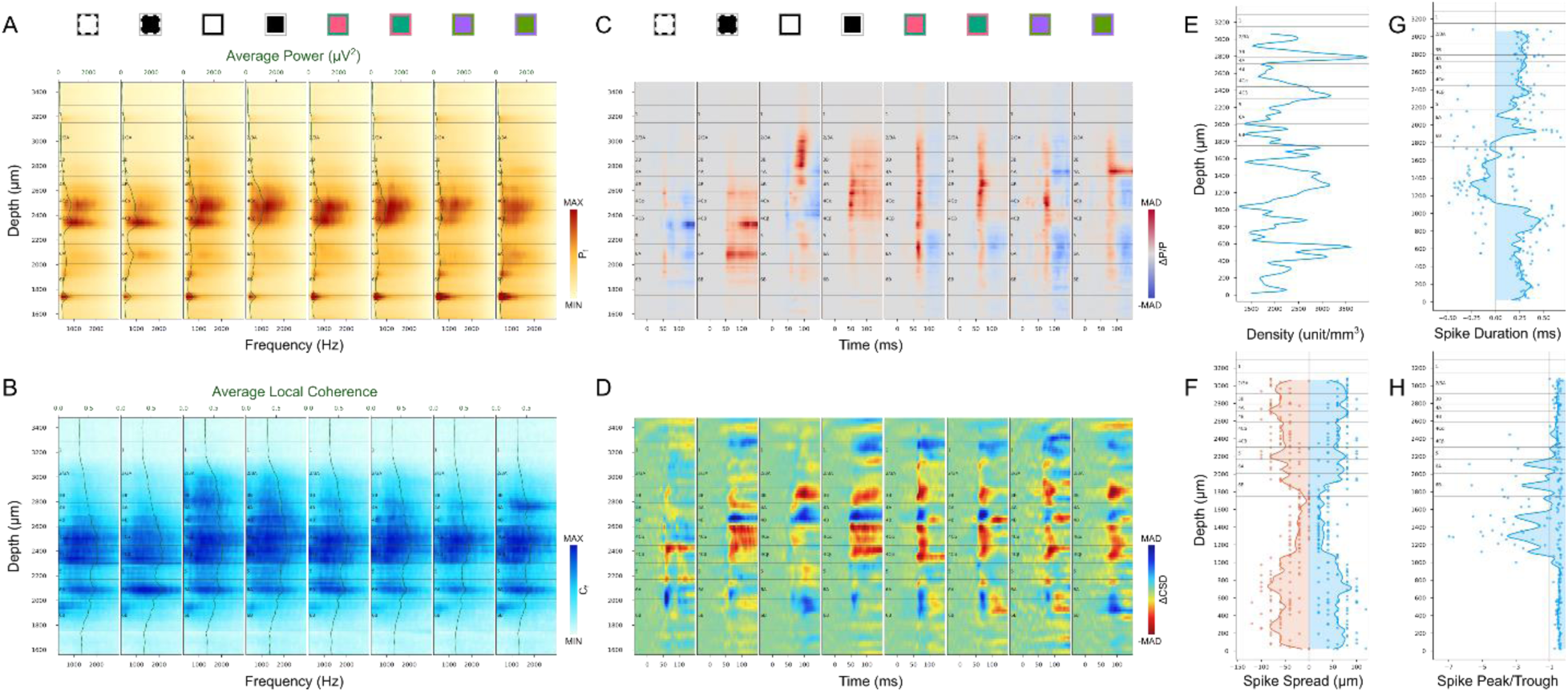
V1 layer delineation for example penetration (B3) based on power spectrum (A), local coherence (B), instantiations power changes (C), CSD changes (D), unit density (E), spike spread (F), spike duration (G) and spike peak-trough ratio (H).

**Figure S7.**
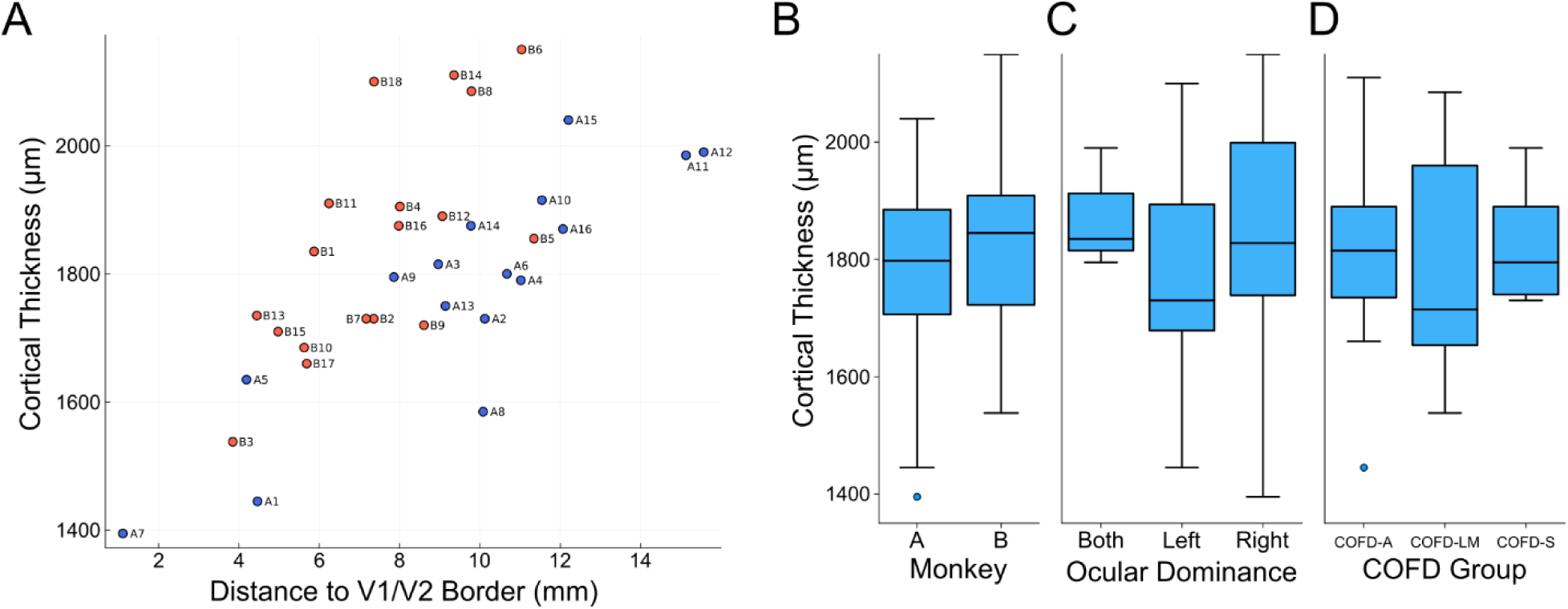
Thickness of primary visual cortex. (**A**) The distance was measured from the location of penetration perpendicular to V1/V2 border. Here, all penetrations except A17 (penetration location information lost) were included. (**B-D**) Distributions of cortical thickness across two monkeys, different ocular dominance columns, and COFD groups. No significant differences were found in each category (Kruskal-Wallis Test, p>0.05). COFD-A: Achromatic ON/OFF domains; COFD-LM: L- and M-cone ON/OFF domains; COFD-S: S-cone ON/OFF domains.

**Figure S8.**
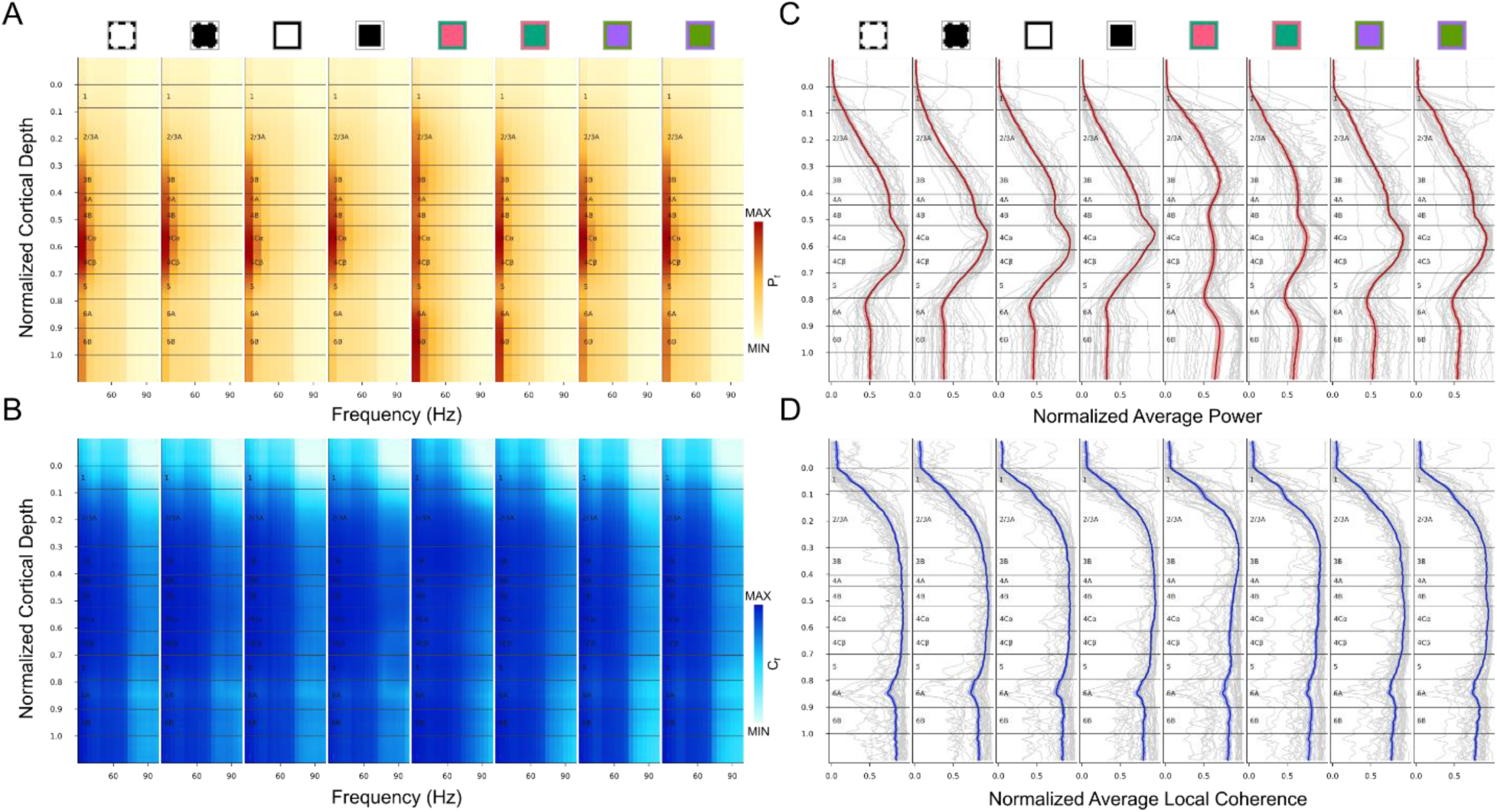
Average metrics from gamma band LFP in V1 template. (**A-B**) Average gamma band (30-100Hz) spectrum profile of power (A) and local coherence (B) across penetrations. The square markers and color bars were the same as the figures in this article. (**C-D**) Normalized average power and local coherence in the gamma band. Gray lines are for each penetration, color-lines and corresponding shaded ribbons show Mean±SEM.

**Figure S9.**
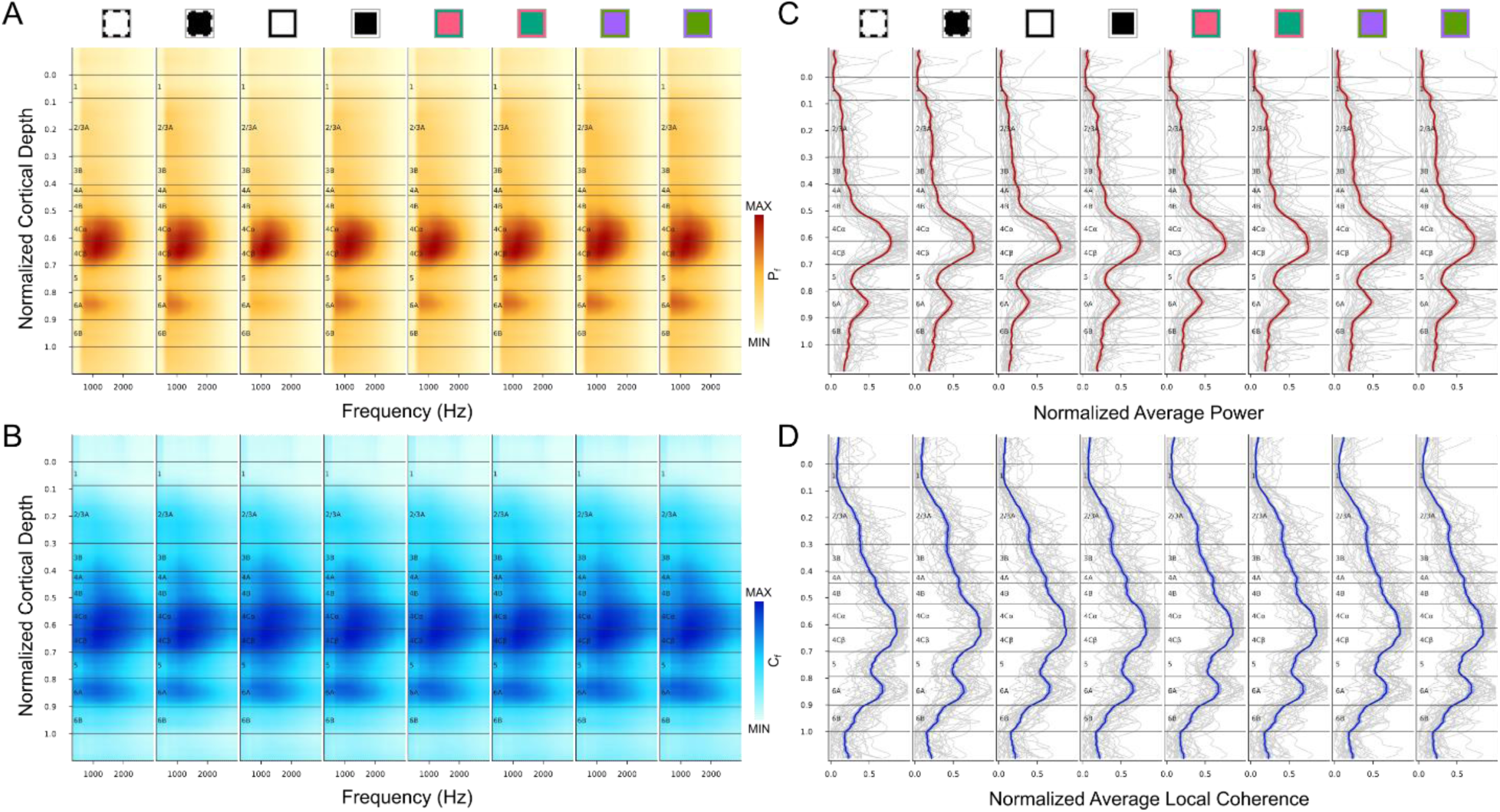
Average metrics from baseline window in V1 template. (**A-B**) Average baseline AP spectrum profile of power (A) and local coherence (B) across penetrations. The square markers and color bars were the same as the figures in this article. (**C-D**) Normalized average baseline AP power and local coherence. Gray lines are for each penetration, color-lines and corresponding shaded ribbons show Mean±SEM.

**Figure S10.**
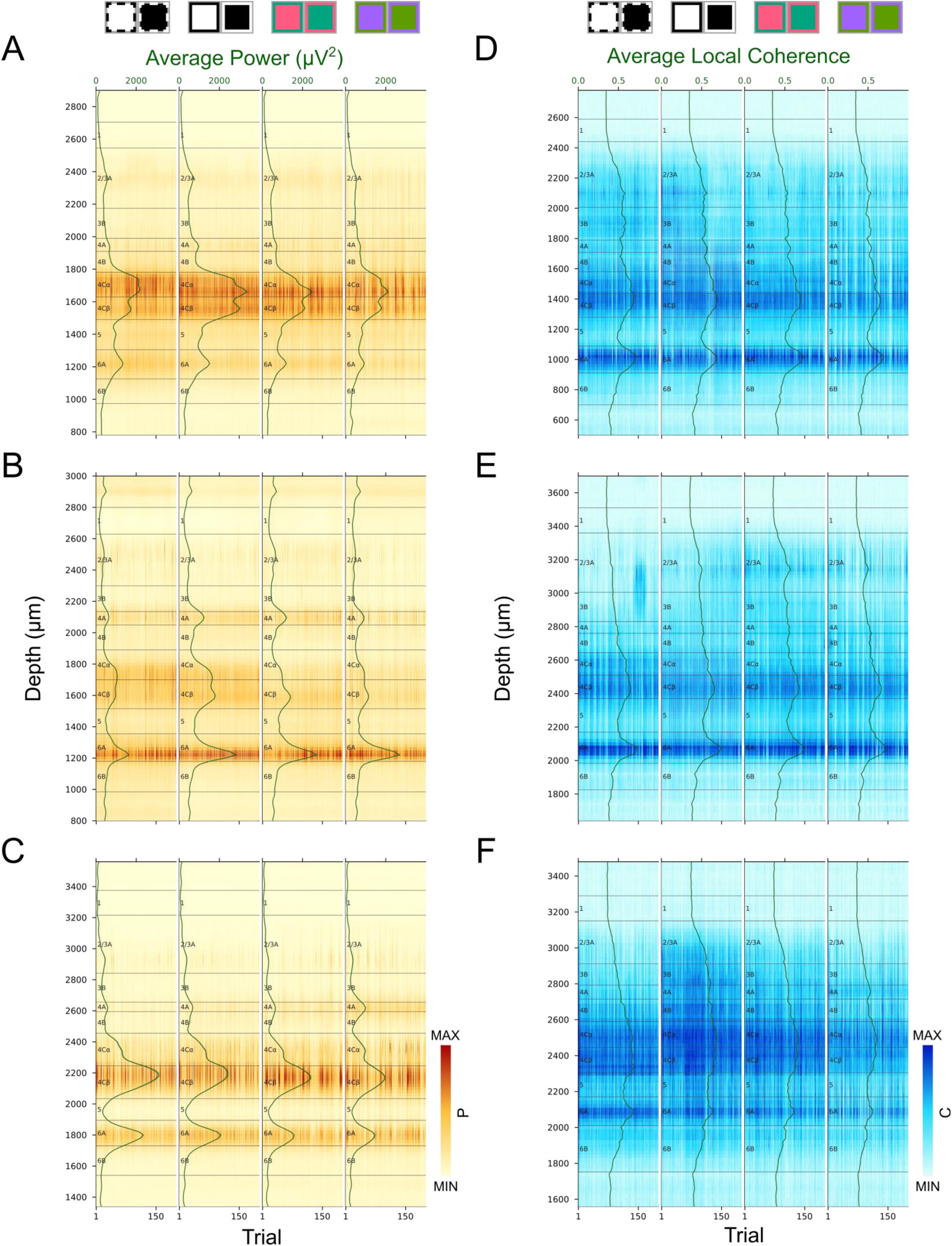
AP power and local coherence averaged across frequencies for each trial. (**A-C**) Powers for the same penetrations (A: A2, B: A3, C: B1) in Figure 5. (**D-F**) Local coherences for the same penetrations (D: B12, E: B10, F: B3) in Figure 6. Power and local coherence profiles were independently color-mapped according to the minimum and maximum in each profile. The green lines and corresponding shaded ribbons showed the Mean±SEM across trials.

**Figure S11.**
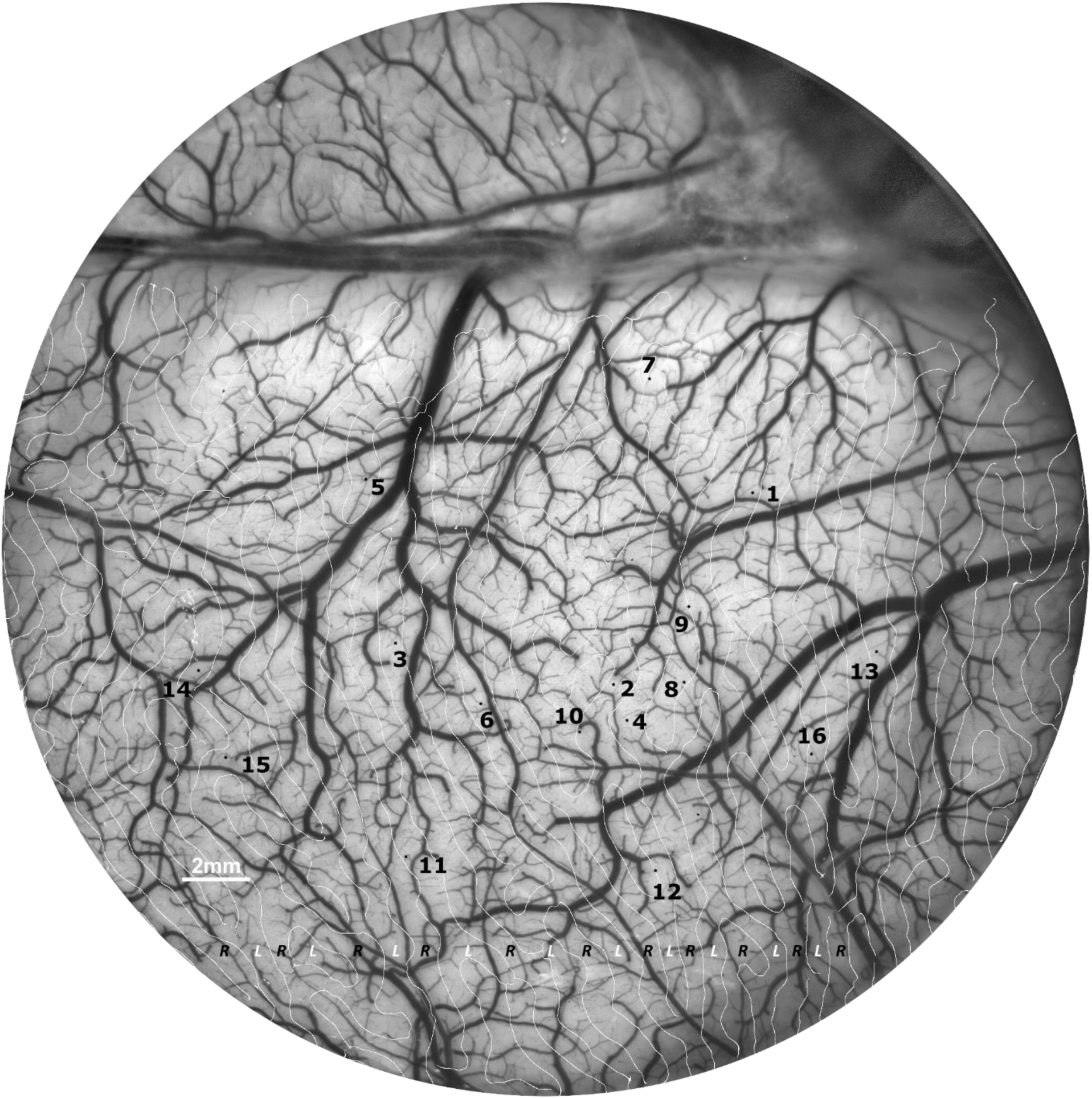
Penetration sites on V1 surface of monkey A. White lines together with L/R texts indicate the borders of ocular dominance column.

**Figure S12.**
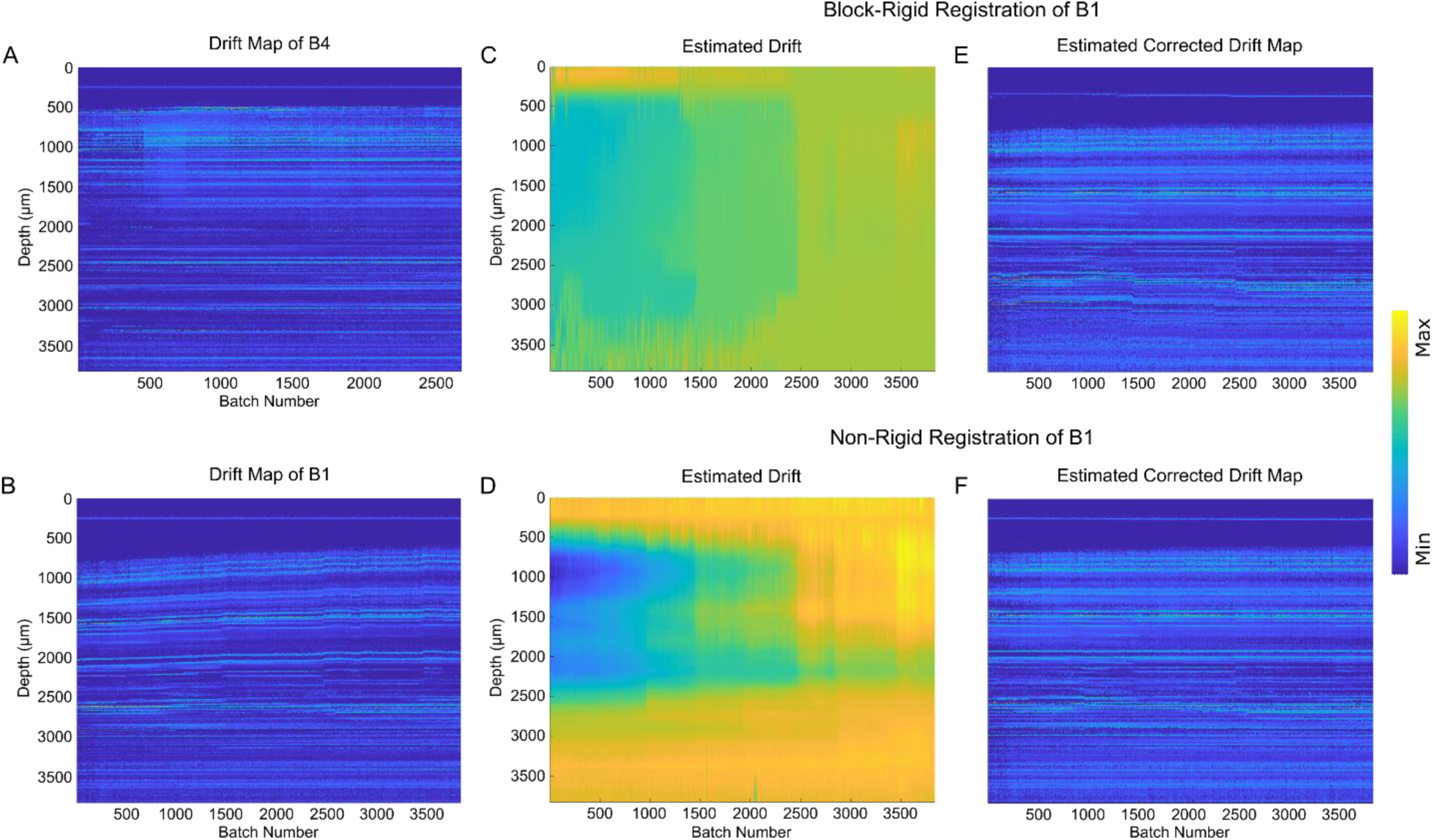
Electrodes drift correction. (**A**) Probe drifting during the recording time for penetration B4 was visualized as average spike amplitudes shifting in consecutive batches (batch length: ∼3.3sec). (**B**) Probe drifting of penetration B1 with same batch length as penetration B4. (**C**) Drift estimated for penetration B1 using the default method in Kilosort 3 (drift range: [-300, 300] µm, number of blocks: 13, drift range for each block: [-300, 300] µm). (**D**) Drift estimated for penetration B1 using ‘imregdemons’ function in MATLAB. (E) Expected correction result of penetration B1 after applying (‘imwarp’ function in MATLAB) the estimated drift in C. (F) Expected correction result of penetration B1 after applying (‘imwarp’ function in MATLAB) the estimated drift in D.

